# Contrasting patterns of somatic mutations in neurons and glia reveal differential predisposition to disease in the aging human brain

**DOI:** 10.1101/2023.01.14.523958

**Authors:** Javier Ganz, Lovelace J. Luquette, Sara Bizzotto, Craig L. Bohrson, Hu Jin, Michael B. Miller, Zinan Zhou, Alon Galor, Peter J. Park, Christopher A. Walsh

## Abstract

Characterizing the mechanisms of somatic mutations in the brain is important for understanding aging and disease, but little is known about the mutational patterns of different cell types. We performed whole-genome sequencing of 71 oligodendrocytes and 51 neurons from neurotypical individuals (0.4 to 104 years old) and identified >67,000 somatic single nucleotide variants (sSNVs) and small insertions and deletions (indels). While both cell types accumulate mutations with age, oligodendrocytes accumulate sSNVs 69% faster than neurons (27/year versus 16/year) whereas indels accumulate 42% slower (1.8/year versus 3.1/year). Correlation with single-cell RNA and chromatin accessibility from the same brains revealed that oligodendrocyte mutations are enriched in inactive genomic regions and are distributed similarly to mutations in brain cancers. In contrast, neuronal mutations are enriched in open, transcriptionally active chromatin. These patterns highlight differences in the mutagenic processes in glia and neurons and suggest cell type-specific, age-related contributions to neurodegeneration and oncogenesis.

## INTRODUCTION

Somatic mutations accumulate in every tissue of the human body throughout life, via mechanisms that depend on intrinsic tissue physiology and exogenous agents^1–8^. Because human tissues comprise diverse cell types with unique properties, quantifying cell type-specific rates and mechanisms of somatic mutation is fundamental to understanding aging and disease initiation at the tissue level. Although previous studies have addressed somatic mutations in aging human neurons^4,9–11^, mutations in glial cells—which represent more than half of the cellular content of the brain and play primary roles in several brain disorders—have not yet been examined. Abnormalities of white matter (WM), which consists mostly of glial cells, are hallmarks of normal brain aging^12,13^ as well as neurodegenerative^14,15^ and neuropsychiatric disorders^16^, but the causes of these changes are unknown. Furthermore, glial progenitor cells are the cell-of-origin of many brain tumors ^17^, and recent findings showed that WM in non-diseased human brain is enriched with clonal oncogenic mutations compared to grey matter^18^.

Oligodendrocytes (OLs) are the main cell type of the WM, and OL dysfunction has been reported in certain brain tumors, age-related disorders ^19,20^, psychiatric disorders ^21,22^ and immune-related multiple sclerosis ^23^. OL generation in humans begins during the second trimester of gestation, peaks at birth and during the first years of life, and continues into adulthood, though at reduced rates^24,25^. Unlike neurons, which mostly arise before birth, OLs are replenished throughout postnatal life by resident oligodendrocyte-precursor cells (OPCs)^24,26^, with the rate of replenishment diminishing with age ^27^. Dysregulation of proliferation and differentiation in the OL lineage is involved in brain cancer, and OPCs are recognized as the cell of origin in some gliomas ^28–31^. Thus, in contrast to neurons, OLs may be subject to mutational processes related to DNA replication and can potentially undergo positive selection relevant for cancer insurgence^32^.

In this study, we assessed genome-wide rates and patterns of aging-related somatic mutations in OLs compared to neurons isolated from the same individuals with single-cell whole-genome sequencing (scWGS). In addition, we generated single-cell Assay for Transposase-Accessible Chromatin with high-throughput sequencing (scATAC-seq) data from these brains and integrated new as well as published^8^ single-cell RNA sequencing (scRNA-seq) data obtained from individuals in the same cohort (Figure 1A). Joint analysis of these data showed OL- and neuron-specific rates and patterns of somatic mutation accumulation, with DNA replication and transcription playing significant roles in OL and neuronal mutagenesis, respectively, and captured features of mutational processes in the differentiated OLs as well as in precursor OPCs. The substantial differences in somatic mutation localization between these two adjacent and interacting cell types informs cell type-specific contributions to distinct age-related diseases.

**Figure 1.**
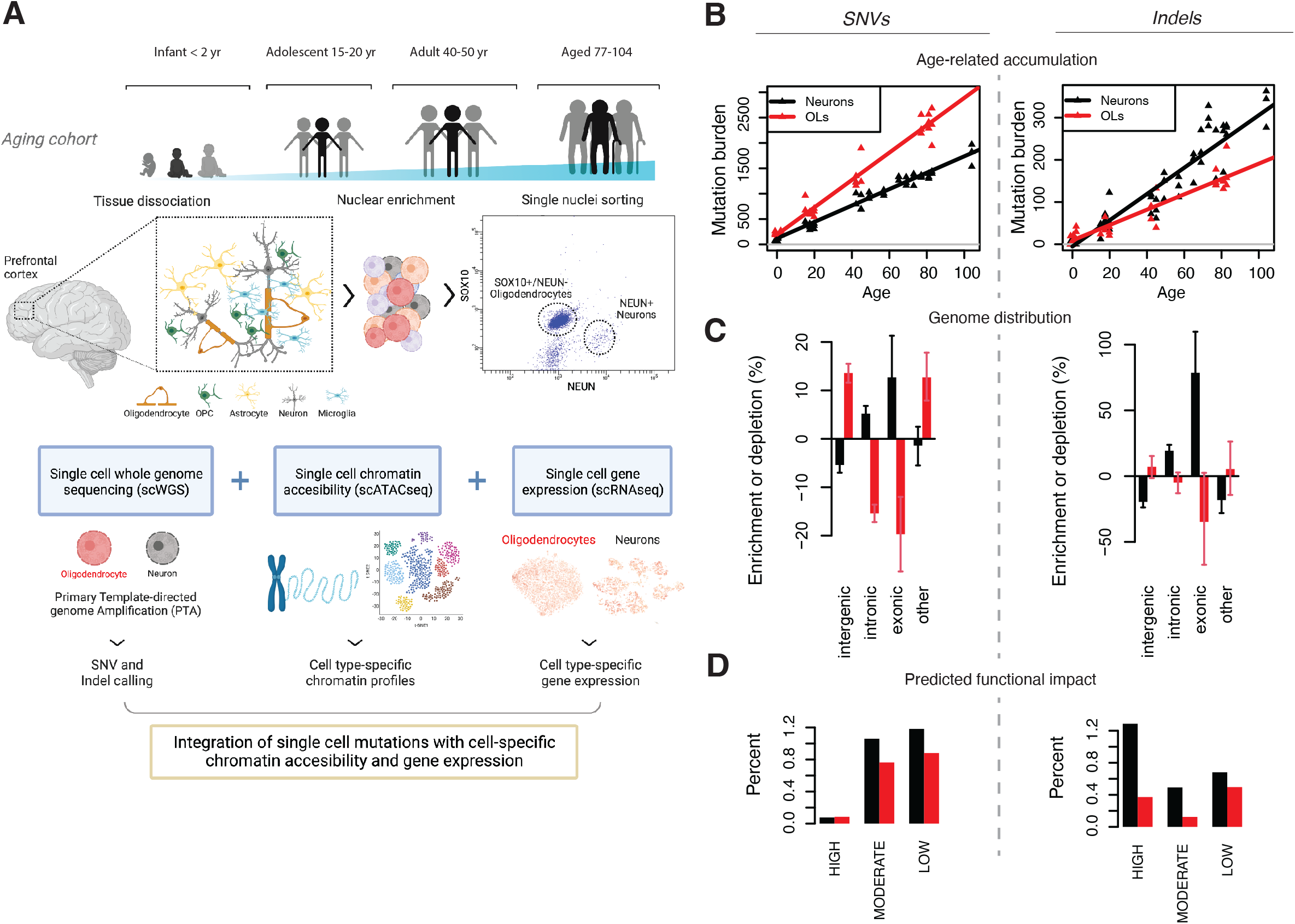
Oligodendrocytes and neurons exhibit contrasting patterns of somatic mutation accumulation. **(A)** Experimental strategy. Oligodendrocytes (OL; n=31 PTA, n=40 MDA) and neurons (n=51 PTA) were obtained from the brains of 17 neurotypical individuals (0-104 years of age) through FANS using NEUN (neurons) and SOX10 (OL) antibodies. Single genomes were amplified using PTA or MDA and non-clonal sSNVs and indels were called using SCAN2. Mutation distributions were compared with scATAC-seq and scRNA-seq data obtained from a subset of the 17 individuals. (**B)** Extrapolated genome-wide sSNV and indel burdens for OLs and neurons as a function of age. Trend lines are mixed-effects linear regression models (see **Methods**). (**C)** Distribution of OL and neuronal sSNVs and indels in genic and intergenic regions. Enrichment/depletion levels are calculated by comparison to a null distribution obtained by randomly shuffling mutations across the genome (see **Methods**). (**D)** Percent of somatic mutations with HIGH, MODERATE and LOW impact on genes, as determined by SnpEff. See also Figure S1 and Figure S2.

## RESULTS

### OLs accumulate somatic mutations at different rates than neurons

OLs were isolated by antibody staining of nuclei prepared from post-mortem cortical brain tissue, selecting SOX10-positive and NEUN-negative nuclei by fluorescence-activated nuclear sorting (FANS). scRNA-seq performed on the sorted populations confirmed >99% purity for both mature OLs and neurons sorted by NEUN positivity (Figure S1, **Methods**). Overall, 71 OLs were obtained from the prefrontal cortex (PFC) of 12 neurologically normal individuals spanning 0.4 to 83 years of age (Table S1); 31 single-OL genomes were amplified by primary template-directed amplification (PTA)^10,33^ and 40 were amplified by multiple-displacement amplification (MDA) before PTA became available. Finally, we combined our single-OL scWGS with 51 PTA-amplified neurons previously generated^10^ from 17 individuals, including 11 which overlap our OL cohort. Notably, due to the higher rate of technical artifacts caused by MDA^10^, we focused on PTA-amplified samples except where indicated. Following scWGS, somatic single-nucleotide variants (sSNVs) and small (1-30 base pair (bp)) insertions/deletions (indels) were identified genome-wide using SCAN2, an algorithm we developed recently^10^ (Table S2). To focus on somatic mutations acquired during aging rather than development, high allele frequency clonal sSNVs and indels were excluded by removing somatic calls supported by one or more reads in matched 30-45X bulk DNA sequencing.

Compared to neurons, scWGS of OLs revealed higher yearly rates of sSNV accumulation but lower rates of indel accumulation. As is the case with neurons and many other cell types^3,4,9–11,34^, the increase in OL sSNV burdens was remarkably linear with respect to age, with a rate of 27 sSNVs/year (95% CI: 25.0-28.9), which is significantly greater than the neuronal rate of 16 sSNVs/year (CI: 14.9-17.4, Figure 1B; for the difference, *P* = 9.6 x 10^-17^, *t*-test for coefficients in a linear mixed model (LMM *t*-test), see **Methods**). At birth, OLs contained 60% more sSNVs per genome compared to neurons (intercept: 191 *vs* 119), though this difference was not significant (*P* = 0.15, LMM *t*-test). Unlike sSNVs, indels accumulated more slowly in OLs than in neurons (1.8 (CI: 1.56-2.07) versus 3.1 (CI: 2.61-3.61) indels/year, respectively, *P* = 1.42 x 10^-5^, LMM *t*-test, Figure 1B). Indel burdens at birth were comparable between cell types. Deletions were more prevalent than insertions in both cell types, consistent with previous reports^10,35^ (Figure S2A); however, OL indels were mostly single-bp deletions, while neurons carried greater numbers of 2-4bp deletions and 1bp insertions (Figure S2B), likely representing distinct mechanisms of indel generation.

OL and neuronal mutations showed opposite biases for genic regions, suggesting different mechanisms of mutagenesis and different consequences for gene integrity. OL sSNVs were enriched in intergenic regions (13%, *P* < 10^-4^; all *P*-values for enrichment analyses based on permutation tests, see **Methods**) and depleted in introns and coding regions (15.4%, *P* < 10^-4^ and 19.7%, *P* = 0.0024, respectively) (Figure 1C). This pattern was replicated in MDA-amplified OLs (Figure S2C, see **Methods**). Neuronal sSNVs were instead overrepresented in genes (12.7%, *P* = 0.0047 in exons; 5.2%, *P* < 10^-4^ in introns) and depleted in intergenic regions (5.4%, *P* < 10^-4^). Indels mirrored these patterns but often with greater effect sizes: in genes, OL indels were depleted (34.8%, *P* = 0.0656 in exons; 4.9%, *P* = 0.17 in introns) whereas neuronal indels were strongly enriched, especially in exons (78.6%, *P* < 10^-4^ in exons; 19.3%, *P* < 10^-4^ in introns), as previously reported^10^. In general, a larger fraction of neuronal mutations were predicted (by SnpEff^36^, see **Methods**) to functionally impact genes (Figure 1D). Strikingly, the rate of indels with the most severe gene-altering effects was ~3-fold higher in neurons than in OLs. Due to the small number of mutations overall, no significant mutation enrichment or depletion was detected for any individual genes after correction for multiple hypothesis testing (Figure S2D).

### OL somatic mutations produce signatures associated with cell proliferation

Analysis of mutational spectra and signatures indicated shared and cell type-specific mutational mechanisms in OLs and neurons. The spectrum of OL sSNVs closely matched the spectra of highly proliferative hematopoietic stem and progenitor cells (HSPCs, cosine similarity 0.946), ^9,34,37,38^ whereas neurons were only marginally similar to HSPCs (cosine similarity 0.773, Figure 2A). Surprisingly, the OL spectrum was less similar to neurons (cosine similarity 0.887) than to HSPCs, likely reflecting a contribution of cell proliferation to OL somatic mutagenesis.

**Figure 2.**
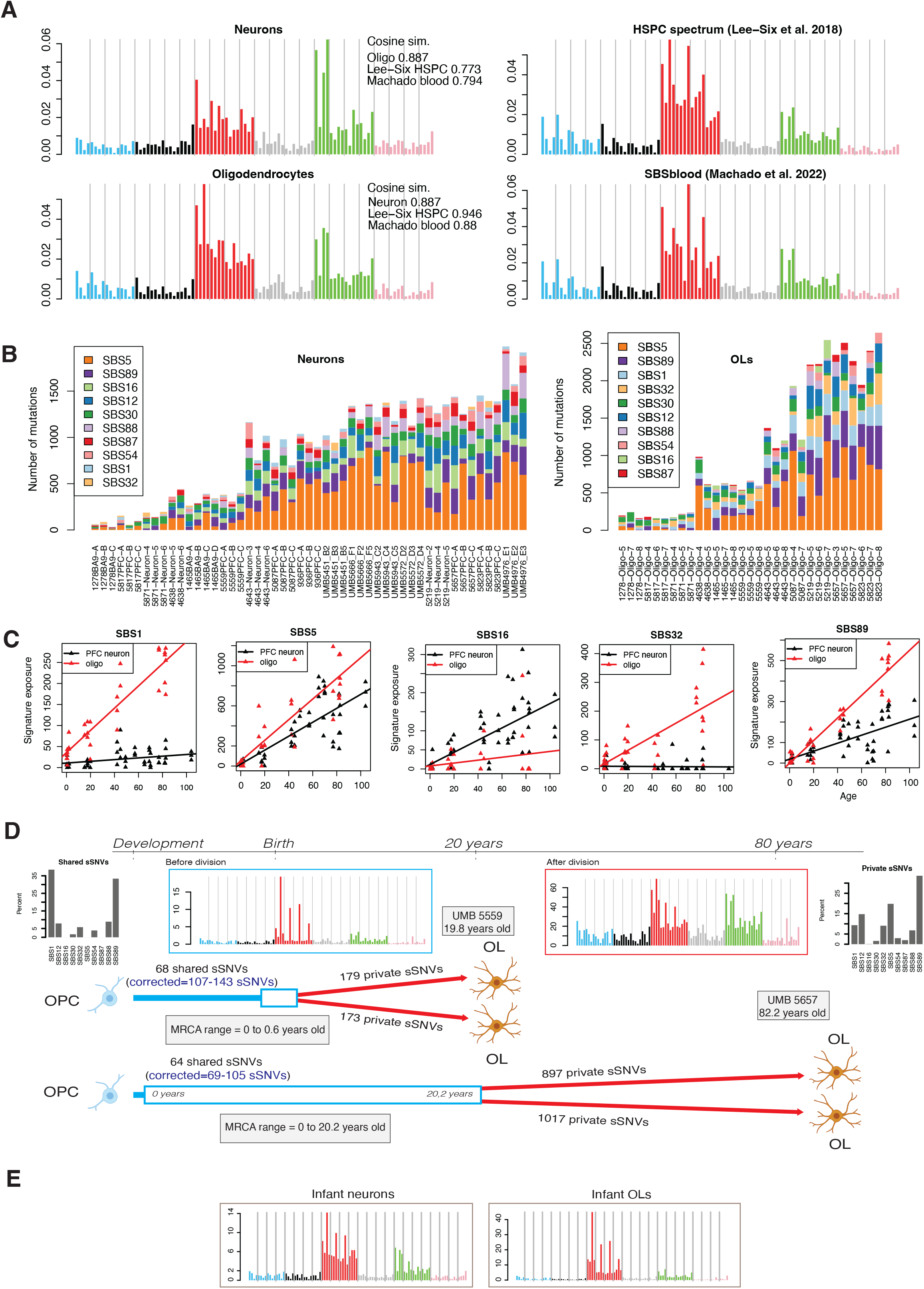
Comparison of single base substitution (SBS) mutational signatures in human oligodendrocytes and neurons. **(A)** SBS mutational spectra of neuronal and oligodendrocyte sSNVs identified in this study (left column); the spectrum of hematopoietic stem and progenitor cells (HSPCs) identified in Lee-Six et al.^37^ and a signature derived from an analysis of human lymphocytes (Machado et al.^38^). Cosine similarities are shown for each pair of spectra. **(B)** The number of somatic mutations, after extrapolation to genome-wide burdens, attributed to each COSMIC SBS signature by least squares fitting for each PTA single OL and neuron. Subjects are ordered from young (left) to elderly (right). A reduced COSMIC catalog of 10 signatures with evidence of activity in either OLs or neurons was used to minimize overfitting (see **Methods**). **(C)** Same as **(B)**, but plotted against age. Each point represents one single cell. Trend lines are mixed-effects linear regression models (see **Methods**). **(D)** Schematic of two pairs of related OLs and estimates of the time of division for each pair’s most recent common ancestor (MRCA). Insets show the SBS spectrum and contributions of COSMIC signatures for sSNVs acquired before division of the MRCA (shared, top left) and sSNVs acquired after division of the MRCA (private, top right). Spectra in panels (D) and (E) are reconstructions obtained by summing exposure-weighted signatures after fitting the infant mutations to the COSMIC SBS catalog (see **Methods**). **(E)** Mutational spectra of all mutations from infant (0-2 years of age) subjects in PTA neurons (left) and PTA OLs (right). The similarity in spectra between infant OL mutations and shared OL mutations supports the estimated time of most recent common ancestor (MRCA) division.

To explore mutagenic mechanisms, we identified single base substitution (SBS) mutational signatures for the identified sSNVs using the COSMIC catalog (v3.1)^39^. Ten of the COSMIC SBS signatures passed our thresholds for activity in either OLs or neurons (Figure 2B, **Methods**) and the remainder were removed to reduce overfitting. Signatures SBS5 and SBS89 were the most prevalent signatures in both cell types, and both signatures accumulated at higher rates in OLs compared to neurons (SBS5, 10.2 versus 6.95 sSNVs/year, *P* = 0.014; SBS89, 5.5 versus 1.95, *P* = 6×10^-11^, Figure 2B-C, LMM *t*-test). SBS5 is a clock-like signature that accumulates independently of cell division, whereas SBS89 was recently reported in colon crypts^3^ but has no known etiology. Signatures SBS1 and SBS32 were strongly associated with age in OLs (*P* < 10^4^ for both signatures, LMM *t*-test) but were nearly absent in neurons (*P* = 0.17 and *P* = 0.55, LMM *t*-test). SBS32 is a C>T signature recently reported as a component of the HSPC spectrum^34^ and SBS1 is a clock-like signature associated with cell division^40^ and accumulated at rates of 2.63 sSNVs/year and 0.20 sSNVs/year in OLs and neurons, respectively (Figure 2B-C). Since mature OLs are post-mitotic, SBS1 may be generated primarily during the OPC stage; if true, the linear accumulation of SBS1 with age in mature OLs implies surprisingly constant levels of OPC division during adult life. Only SBS16, a signature associated with transcription, accumulated at a higher rate in neurons, with 1.59 sSNVs/year compared to 0.38 sSNVs/year in OLs, consistent with the enrichment of neuronal mutations in transcribed genomic regions and in line with previous reports^4,10^.

Two pairs of closely related OLs, which likely trace their ancestry to common OPCs, illustrate the extent to which signatures of early- and late-life mutations differ, and represent permanent maps of developmental lineages and age-related changes. Despite filtering high allele frequency clonal sSNVs by removing mutations found in bulk tissues, two OL pairs from two individuals (subjects 5559 (PTA OLs) and 5657 (MDA OLs), 19.8y and 82y, respectively) shared unusually high levels of sSNVs (68 and 64 sSNVs, respectively, Figure 2D), indicating common ancestry. In both subjects, the shared sSNVs were mostly C>T transitions at CpG sites and fitting to the COSMIC signatures revealed a 38% contribution from the cell-division-related signature SBS1. Each pair of OLs also contained similar numbers of private sSNVs (179 and 173 for subject 5559; 897 and 1017 for subject 5657), consistent with equal lifetimes for each of the cognate OLs after the division of their most recent common ancestor (MRCA) OPC. The mutational spectrum of private sSNVs was similar to the OL spectrum (Figure 2A,D) and, after fitting to COSMIC and removing MDA-associated artifacts (**Methods**), the spectrum was primarily explained by SBS89 and SBS5 (34% and 20%), followed by SBS12, SBS1 and SBS32 (15%, 9% and 9%, respectively, Figure 2D). We estimated the age at which the MRCA divided in each subject by adjusting private sSNV counts for calling sensitivity and dividing by our yearly sSNV accumulation rate (see **Methods**). This placed the MRCA of subject 5559 near birth and the MRCA of subject 5657 between 0-20 years of age. Comparison of mutation spectra with infant OLs (subjects aged 0-2 years old) provided orthogonal evidence that the MRCA occurred during infancy. Indeed, the spectrum of shared sSNVs matched the spectrum of infant OL sSNVs with a cosine similarity of 0.97 (Figure 2E). Crucially, the neuronal sSNV spectrum from the same infant subjects contained far fewer SBS1-like C>Ts at CpG dinucleotides, implying that the increase in SBS1 occurred in an OL-specific lineage and does not reflect early clonal sSNVs that evaded our filters. We speculate that the most likely MRCA time coincides with a burst of OL generation that occurs in the young human brain (0-10 years of age)^24^. The relationships of these two pairs of cells suggest that shared mutations mark a permanent forensic lineage tree, while non-shared mutations represent a linear timer of when any two cells separate from a common progenitor.

Indel (ID) signatures revealed shared and specific mutational processes further distinguishing OLs from neurons (Figure 3A-C). ID4, a signature representing ≥2bp deletions and associated with transcriptional mutagenesis^41^, was most strongly correlated with age in neurons, as previously reported ^9,10^ but was almost completely absent in OLs (*P* = 0.534, LMM *t*-test; Figure 3C). ID5 and ID8, two clock-like indel signatures, were present in both cell types, with ID8 correlated more strongly with age in neurons than in OLs. The two remaining clock-like indel COSMIC signatures, ID1 and ID2, were not detected, but they are difficult to identify in PTA data due to similarity with sequencing artifacts^10^. ID9, which is characterized by 1bp deletions, was the most prevalent signature in OLs and accumulated at a rate of 0.54 indels/year; in neurons, the accumulation was significantly lower at 0.20 indels/year (*P* = 0.0017, LMM *t*-test). Interestingly, this ID9 signature is also found in a large fraction of adult gliomas^42^ as well as in a considerable fraction of other brain tumors^39^.

**Figure 3.**
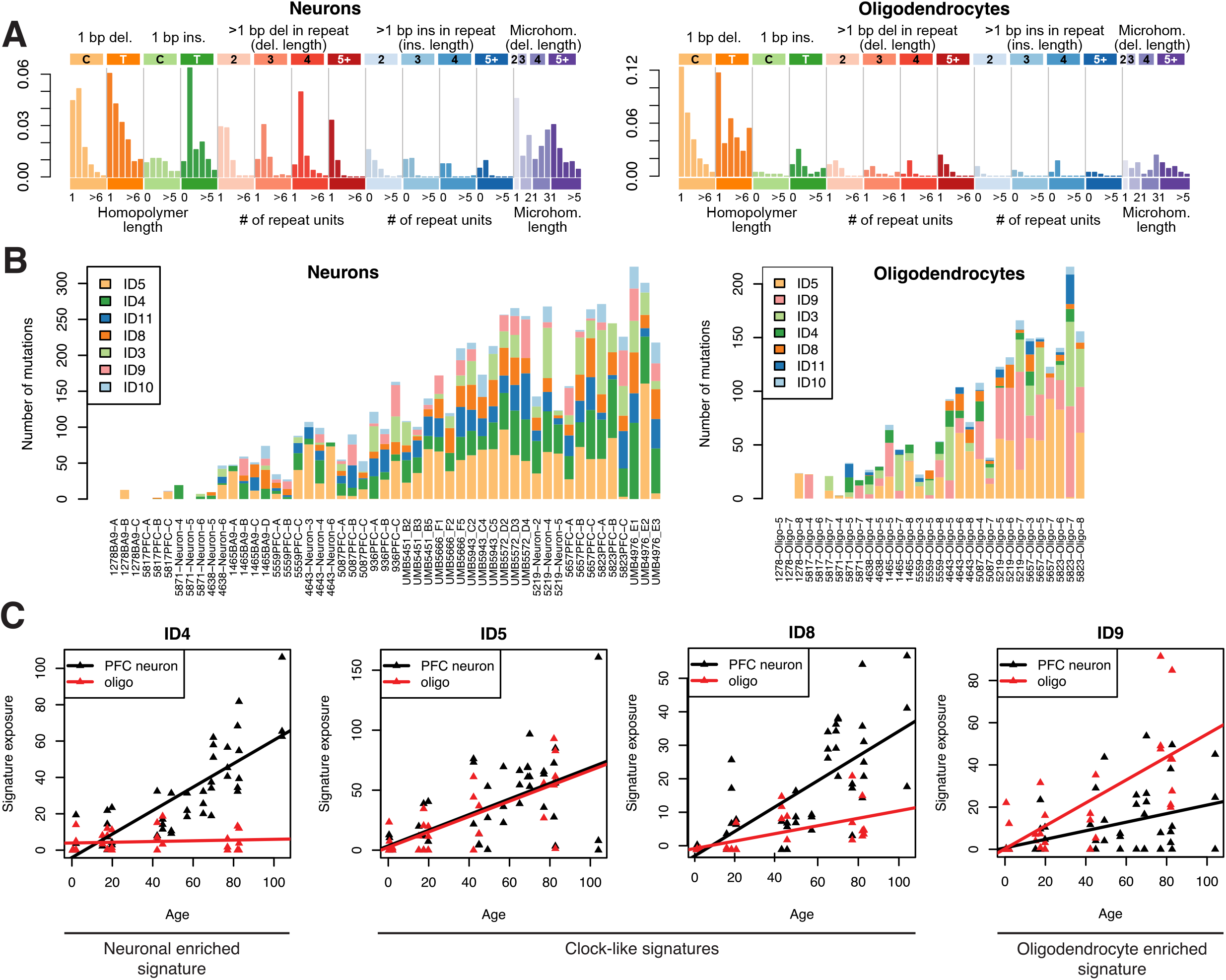
Comparison of insertion and deletion (ID) COSMIC signatures in human oligodendrocytes and neurons. **(A)** Spectra of somatic indels from human OLs and neurons using the 83-dimensional indel classification scheme from COSMIC. **(B)** Contribution of COSMIC indel signatures to each single OL and neuron. One bar represents one single cell; cells are ordered according to age with the youngest individuals on the left and eldest individuals on the right. **(C)** Same as **(B)**, but signature exposure is plotted against age for each single cell; each point represents one cell. Trend lines are mixed-effects linear regression models (see **Methods**). ID5 and ID8 are annotated as clock-like signatures in COSMIC.

### OL SNVs are enriched in inactive genomic regions

Our earlier observation that OL mutations were depleted in genes—opposite to the pattern in neurons (Figure 1C)—suggested different determinants of mutagenesis in these two cell types. Comparison of somatic mutation density to additional data types, including scRNA-seq, scATAC-seq, replication timing, and chromatin marks, revealed that OL mutations are enriched in chromatin that is either inaccessible, untranscribed, or which harbors repressive histone marks—which we refer to as inactive chromatin—in striking contrast to neuronal mutations. We first compared somatic mutation densities to gene expression levels from brain scRNA-seq data for three subjects in our cohort (1465, 4638 and 4643; 40,083 PFC cells in total; Figure 4A) ^8^. OL sSNVs were depleted by 25%-32% in the top few deciles of expression measured in OLs (Figure 4B, *P* < 10^-4^; all *P*-values in this section are from permutation tests) and were similarly depleted in all other cell types detected in our scRNA-seq data. The negative association between transcription level and somatic mutation density in OLs was confirmed using bulk brain RNA-seq data from the Genotype Tissue Expression Consortium (GTEx)^43^ (Figure S3A). Indels in OLs were not significantly enriched or depleted, possibly due to a lack of statistical power caused by the relatively low number of somatic indels in OLs (Figure 4B).

**Figure 4.**
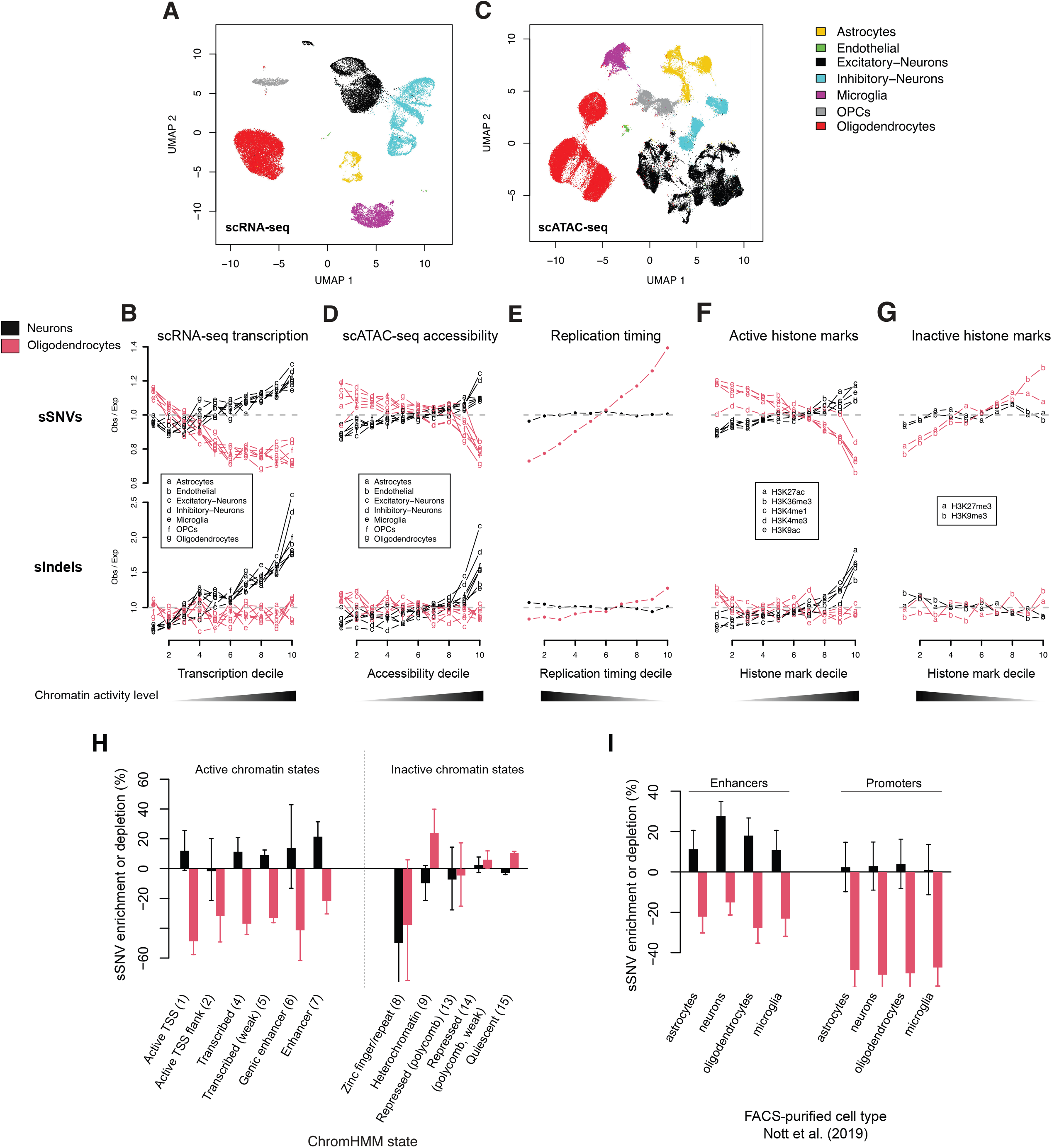
Oligodendrocyte somatic mutations are associated with inactive chromatin while neuronal mutations associate with active chromatin. **(A)** Representative UMAP plot of scRNA-seq from one normal brain from our cohort (subject 1465) with cell type annotations. **(B)** Enrichment analysis of somatic mutations vs. scRNA-seq transcription level. The genome is divided into 1 kilobase, non-overlapping windows and each window is annotated with an average gene expression level per cell type; windows that are <80% covered by a gene are discarded. The remaining windows are classified into 10 deciles, with 1 representing the least transcribed and 10 representing the most transcribed. In each decile, the observed number of somatic SNVs and indels is compared to a null distribution of mutations obtained by randomly shuffling mutation positions (see **Methods**). Enrichment analyses show somatic mutation density vs. transcription level from each of the cell types identified in our scRNA-seq. **(C-D)** Same as (**A**) for scATAC-seq from the brains of 10 subjects from this cohort. **(E)** Somatic mutation density vs. replication timing as measured by ENCODE RepliSeq; lines represent average enrichment across 15 cell lines. **(F-G)** Somatic mutation density vs. 5 epigenetic marks related to gene activity (**F**) and two repressive epigenetic marks (**G**) measured in dorsolateral prefrontal cortex tissue (Roadmap Epigenomic Project, reference epigenome E073). **(H)** Enrichment of somatic mutations vs. functional genomic regions identified by ChromHMM in reference epigenome E073. Numbers in parenthesis indicate the ChromHMM state number. (**I**) Enrichment of somatic mutations vs. active enhancers and promoters identified in Nott et al.^47^ for several brain cell types. See also Figure S3.

Next, brain scATAC-seq data representing ~82,000 cortical cells obtained from ten subjects in our cohort (see **Methods**) revealed a strong depletion of OL sSNVs in open chromatin (Figure 4C,D). In the decile of the genome with the highest chromatin accessibility from OLs identified in scATAC-seq data, OL sSNVs were depleted by 29% (*P* < 10^-4^). Slightly weaker sSNV depletions were observed for the remaining cell types (mean 21% for the top decile of chromatin accessibility), with OPCs showing the second strongest depletion signal (24%, *P* < 10^-4^). A weak but negative trend between OL indel density and chromatin accessibility was also observed (Figure 4D).

Data from the Encyclopedia of DNA Elements (ENCODE) ^44^ and the Roadmap Epigenomics Project^45^ further confirmed enrichment of OL mutations in inactive chromatin. First, OL sSNVs were significantly enriched in late-replicating regions of the genome (which tend to be less transcriptionally active) as determined by RepliSeq data from the ENCODE project (mean 39% in the latest replicated decile, *P* < 10^-4^; Figure 4E, S3B). Comparison to histone marks from the Roadmap Epigenomics Project revealed negative associations between OL sSNVs and marks of open chromatin, transcription and active regulatory elements (H3K27ac, H3K36me3, H3K4me1, H3K4me3 and H3K9ac) and positive associations with repressive marks (H3K9me3 and H3K27me3)^46^ (Figure 4F-G). Chromatin state annotations from ChromHMM^46^, which classify chromatin based on an ensemble of histone marks, further confirmed the pattern of OL mutation enrichment in inactive or inaccessible genomic regions, with OL sSNVs overrepresented in heterochromatin (state 9, 24% enrichment, *P* = 0.0036) and quiescent regions (state 15, 11% enrichment, *P* < 10^-4^), and depleted in the transcriptionally active states 1-7 (Figure 4H). The strongest depletion of OL sSNVs across all genomic covariates analyzed in this study was observed for active transcription start sites (ChromHMM state 1, mean 50.4% depletion across brain tissues) and was not specific to any of the 13 Roadmap Epigenomics brain tissues (Figure S3C). An orthogonal dataset of active promoters in neurons, oligodendrocytes, microglia and astrocytes from flow-sorted cell populations^47^ further confirmed the strong depletion of OL sSNVs in promoters (mean depletion 51%) and again there was no marked preference for the cell type from which the promoters were measured (Figure 4I).

The distribution of neuronal mutations differed from OLs across all the genomic covariates we tested: neuronal sSNV and indel rates increased with gene expression, chromatin accessibility and active histone modifications and decreased with inactive histone modifications (Figure 4A-G). Unlike OLs, somatic mutations in neurons were more specifically associated with transcription levels measured in brain tissues (Figure S3A) and especially with single cell transcriptomic and chromatin accessibility signals from neurons (Figure 4B,D). Neuronal mutations showed little association with replication timing (Figure 4E), which is unsurprising since most neuronal mutations are acquired in the post-mitotic state and clonal somatic mutations were largely removed by our bulk filters.

To further understand the action of mutational processes in the two cell types, we correlated SBS mutation signature exposures (rather than total mutation density) to the previously discussed genomic covariates. In OLs, SBS1 density generally followed the patterns of total mutation density, with positive associations with inactive chromatin and late replication timing (Figure 5A). The distribution of SBS1 in neurons mimicked that of OLs and, in particular, was strongly positively associated with replication timing, suggesting that neuronal SBS1 may have accumulated during cell divisions in neurogenesis. SBS16 (a T>C signature associated with transcriptional activity) exposure in neurons was positively associated with active histone marks, gene expression, and chromatin accessibility levels from excitatory and inhibitory neurons (Figure 5B). Consistent with the known transcribed-strand bias of SBS16, neuronal T>C mutations exhibited the largest transcribed-strand bias of all mutation types in this study (Figure S4). Interestingly, despite neurons being post-mitotic, SBS16 density trended negatively with replication timing, likely reflecting higher gene density in early replicating regions.

**Figure 5.**
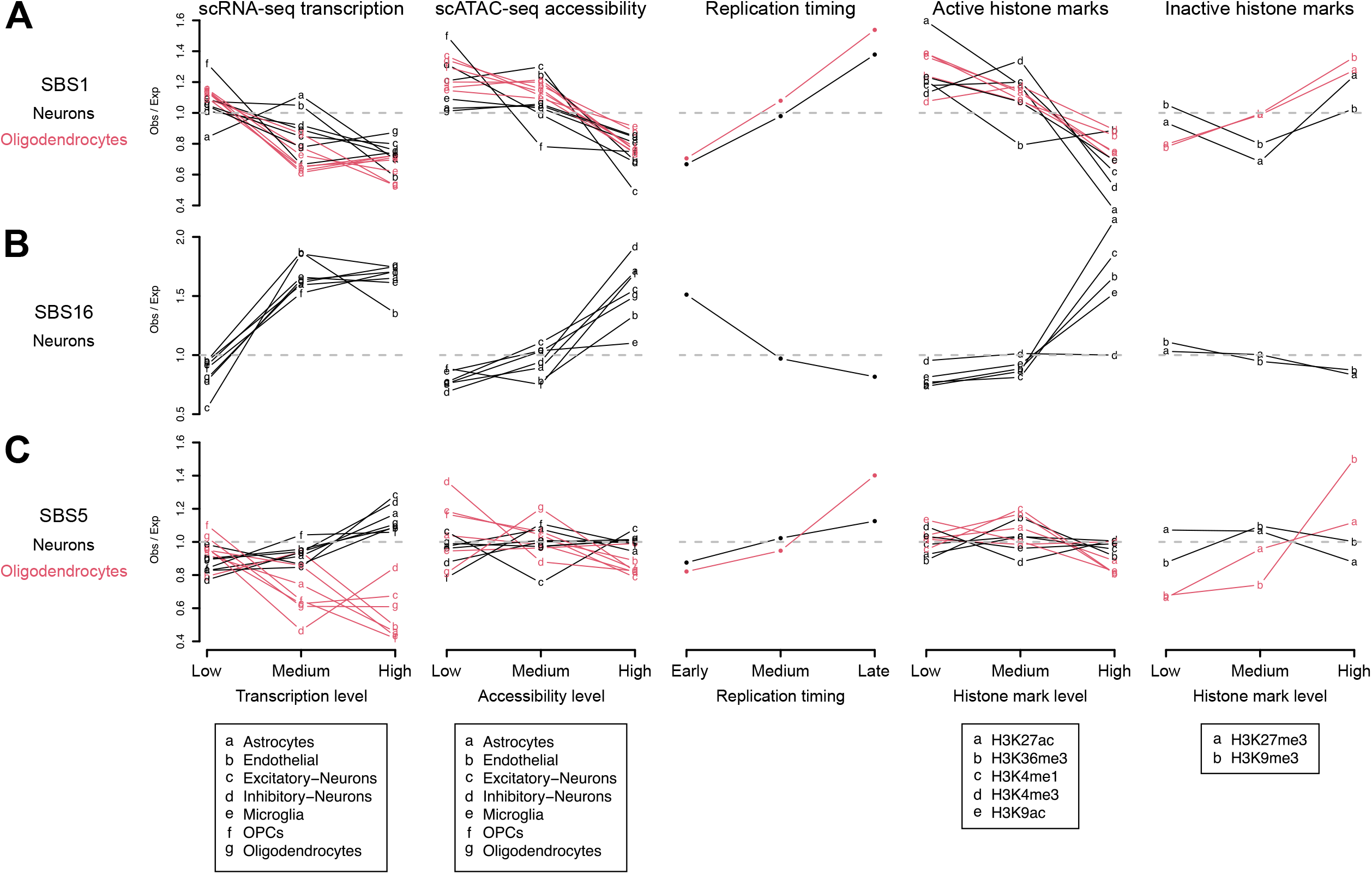
Enrichment of mutational signatures in active and inactive chromatin. Enrichment analysis of somatic mutations attributed to SBS1 **(A)**, SBS16 **(B)** or SBS5 **(C)—** rather than total mutation density—vs. the genomic covariates presented in Figure 4. To reduce noise caused by the smaller number of mutations attributed to each specific mutational signature, the genome has been divided into three quantiles rather than ten. OLs are not plotted for SBS16 due to noisy measurements; none of these omitted OL SBS16 enrichment tests achieved significance at the *P* < 0.05 level. See also Figure S4.

Although SBS5 is the most prevalent signature in both OLs and neurons, it did not accumulate in the same genomic regions in these two cell types, particularly with respect to expression levels (Figure 5C). In OLs, patterns of SBS5 exposure showed little difference from the aggregate somatic mutation density, with negative associations with active epigenetic marks, gene expression and open chromatin and positive associations with inactive marks and late replicating regions. However, in neurons, SBS5 was not significantly associated with any of the covariates tested, with only a marginally significant positive trend with scRNA-seq expression in excitatory neurons (top 33%, *P* = 0.08, Figure 5C). These observations suggest that either SBS5 is generated by cell type-specific mechanisms or that SBS5 may not be a fully decomposed signature—in particular, it may be contaminated by the transcription-associated SBS16, consistent with the marginally significant association with expression levels in neurons—as previously suggested^9^.

### The OL mutation density profile resembles that of glial-derived tumors

Patterns of somatic mutation in cancer often contain sufficient information to identify the cell type from which a tumor emerged^48^; thus, we explored whether our normal OL sSNV densities resembled those from a large collection of cancer WGS data from the Pan-Cancer Analysis of Whole Genomes (PCAWG) project ^42^. OL sSNVs were positively correlated with somatic mutation densities of all cancer types from PCAWG whereas neuronal sSNVs were not correlated with any tumor type (Figure 6A). Specifically, for OL mutations, the highest correlations observed corresponded to glial-derived tumors of the central nervous system (CNS) for which OPCs are thought to be the cell of origin (CNS-GBM, glioblastoma multiforme and CN S-Oligo, oligodendroglioma)^28–31^.

**Figure 6.**
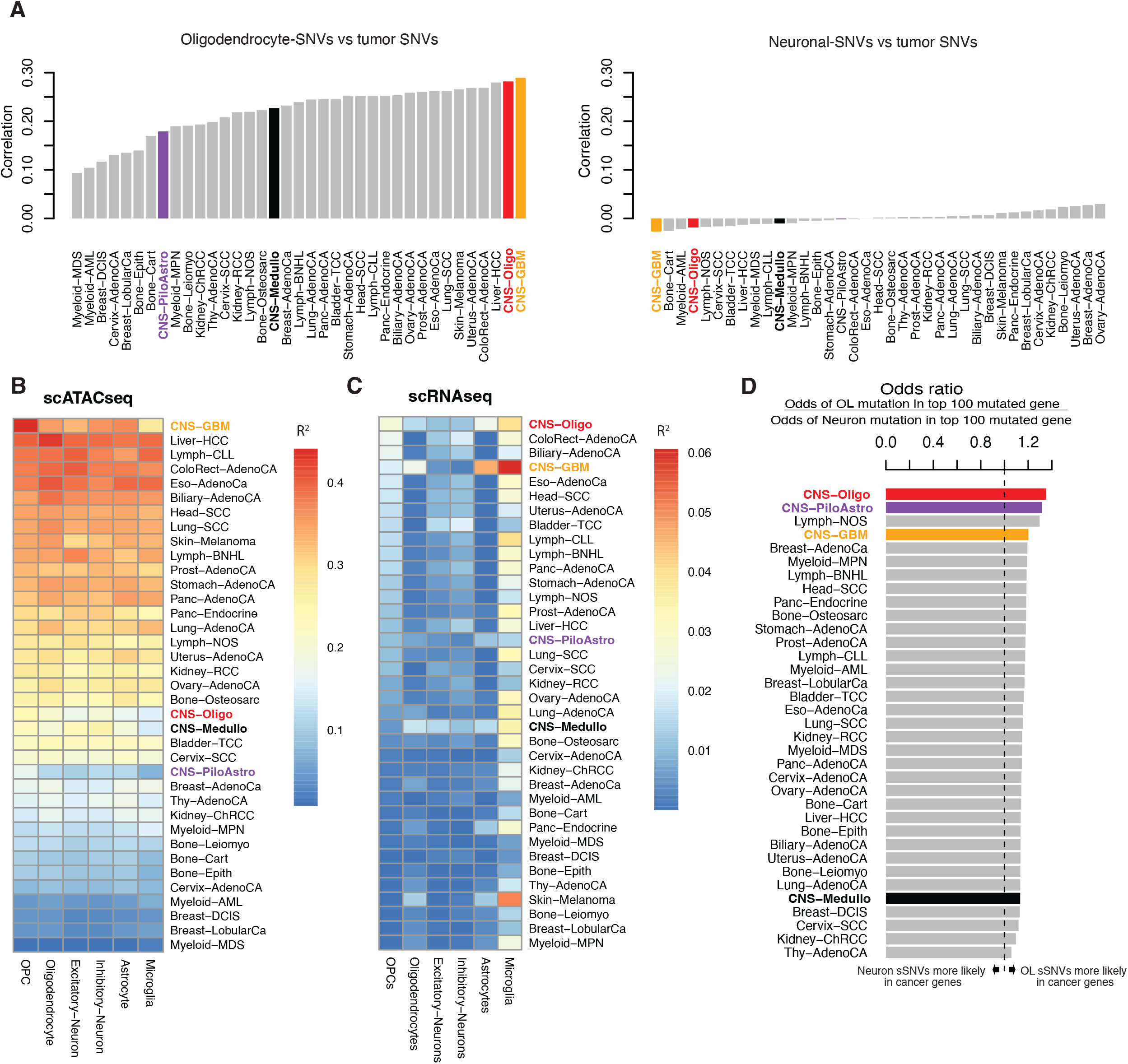
Patterns of oligodendrocyte sSNVs correlate with somatic mutation density in cancer. **(A)** Correlation of OL and neuronal sSNV mutation density with cancer mutation density. For each cell type and cancer type, the genome was tiled with non-overlapping 1 MB bins and numbers of mutations per bin were tabulated. CNS tumors are colored: CNS-Oligo, oligodendroglioma, red; CNS-PiloAstro, pilocytic astrocytoma, purple; CNS-GBM, glioblastoma multiforme, orange; CNS-Medullo, medulloblastoma, black. **(B-C)** Mutation density for each tumor type was fit using a linear regression to cell type-specific single cell chromatin accessibility signals (**B**) and single cell expression levels (**C**) using the same 1 MB bins described in panel (**A**). For each tumor type and cell type, the fraction of variance in tumor mutation density explained (*R*^2^) by each cell type is shown. **(D)** Comparison of OL and neuron somatic mutation rates in frequently mutated cancer genes. For each tumor type in PCAWG (y-axis), the 100 most frequently mutated genes were determined. For each tumor-specific set of 100 cancer genes (*G_T_*) an odds ratio (OR) is computed such that OR > 1 indicates that OL mutations are more likely to occur in *G_T_* and OR < 1 indicates neuronal mutations are more likely to occur in *G_T_*. Formally, OR = (# OL sSNVs in *G_T_* / # genic OL sSNVs not in *G_T_*) / (# neuron sSNVs in *G_T_* / # genic neuron sSNVs not in *G_T_*). See also Figure S5.

Our scATAC-seq data allowed further cell type-specific evaluation of cancer sSNV densities. Among all tumor types in PCAWG, GBM sSNV density was best predicted by OPC-specific scATAC-seq tracks using a regression model, with 44% of variance in GBM sSNV density explained (Figure 6B). This provides additional evidence that OPCs are the cell of origin for GBM tumors and that scATAC-seq is a powerful approach for determining the cell-of-origin for a tumor^48^. Expression levels from scRNA-seq were far less effective in explaining cancer mutation density, explaining only 6% of variance in the best case (Figure 6C).

Finally, we tested whether cancer-associated genes were more likely to be mutated in OLs compared to neurons. For each tumor type, we determined the 100 most frequently mutated genes and computed an odds ratio (OR) indicating whether mutations in OLs (OR > 1), neurons (OR < 1) or neither cell type (OR=1) were more likely to occur in the frequently mutated genes. In general, OL sSNVs were biased toward cancer-associated genes (OR near 1.1 for most cancer types, Figure 6D), likely reflecting the general similarities between OL and cancer sSNV mutation densities. However, OL sSNVs were clearly biased toward genes mutated in CNS tumors, with the highest odds ratios observed for oligodendrogliomas (CNS-Oligo, OR=1.35, *P* = 1.6×10^-7^, Fisher’s exact test) and pilocytic astrocytomas (CNS-PiloAstro, OR=1.31, *P* = 3.1×10^-6^). Analysis of the top *N* cancer mutated gene lists for *N* = 10 to 500 confirmed that these findings did not depend on our choice of cutoff *N* = 100 (Figure S5). Altogether, the similarities between OL and cancer mutation patterns suggest a contributing relationship between OL mutations—especially those acquired at the OPC stage—and tumorigenesis.

## DISCUSSION

Our integrative analysis of somatic mutations uncovered OL-specific mutational processes during aging compared to neurons and suggests how these differences may predispose to diseases such as cancer. Our study design provides a unique opportunity to explore how different cell types sharing the same microenvironment for years—or even decades—can exhibit contrasting features that ultimately shape the human brain. An additional advantage of our design is that comparison of OLs and neurons measured by the same single-cell DNA sequencing technology helps to rule out the possibility that differential mutation rates or genomic distributions reflect technical artifacts or biased representation of specific genomic regions.

Somatic mutation burdens increase linearly in both OLs and neurons with age; however, OLs accumulate 69% more sSNVs than neurons and 42% fewer indels. The apparent low indel rate in OLs may reflect a high rate of indel mutagenesis in neurons compared to other cell types, as reported by previous studies^9^. Some of the excess sSNVs in OLs (e.g., those attributed to SBS1) are likely associated with cell division in ancestral OPCs^49^, but due to lacking information regarding the mechanisms of SBS5, SBS32 and SBS89 mutagenesis, it is not clear what processes account for the remaining excess sSNV burden in OLs. A recent study highlighted the importance of cell proliferation-independent sources of somatic mutations in normal cells and hypothesized that the interplay between cell type-specific DNA damage and repair processes may underlie differences in mutation burden between cell types^9^. Hence, less efficient DNA repair processes in OLs—rather than additional DNA damage—may be a plausible explanation for the excess OL sSNV burden compared to neurons. Follow-up studies mapping DNA repair sites in OLs versus neurons might be needed to address this question^50–52^. Another important factor differentiating mutations in OLs and neurons is the strength of selective forces acting during aging. Aging neurons cannot be subject to positive selection, and negative selection is likely limited to highly deleterious mutations that induce cell death. OLs, on the other hand, include mutations accumulated by postnatal cycling OPCs in which both positive and negative selective effects may be present. Thus, the subset of OL somatic mutations acquired at the ancestral OPC stage are of particular importance since they can expand clonally and amplify deleterious effects.

OL mutations are more prevalent in transcriptionally inactive and/or inaccessible chromatin and resemble patterns reported in cancer^32^ and other proliferative cells^48^. Associations between OL somatic mutation density and genomic covariates generally were not cell type- or tissue-specific; however, it is important to interpret this observation in light of the fact that OL mutations represent a combination of mutations accrued during the OPC and mature OL states. Neuronal mutations were characterized by a strongly contrasting pattern of enrichment in transcriptionally active, open chromatin with clear preference for genomic covariates measured in brain tissue and, specifically, excitatory neurons.

Mutational signature analysis was helpful in interpreting some contributing factors to the overall mutational burden and to its accumulation over time. SBS1 was prevalent in OLs and nearly absent in neurons, consistent with previous characterizations of SBS1 as a cell division-dependent mutational clock^40^ and at odds with a recent study which estimated a nearly 10-fold greater SBS1 rate in human neurons^9^. SBS5 made up the majority of mutations in both OLs and neurons, but was distributed differently across the genome in the two cell types. One attractive explanation for this is differential repair: i.e., that SBS5-associated DNA damage may occur throughout the genome but be more efficiently repaired in certain genomic regions in a cell type-specific manner. However, concerns of contamination by other signatures and other algorithmic issues may yet be to blame; for example, the current COSMIC catalog was generated primarily by cancer exomes and genomes, and signatures present in post-mitotic cells are likely to be under-represented. In addition, despite the dozens of single cells we sequenced, the total number of mutations is not large enough to confidently identify signatures that are present at low exposures.

Our work demonstrates marked differences in somatic mutation accumulation between neurons and OLs in the same tissue and reveals the mutational dynamics of OLs during neurotypical brain aging. In contrast to neuronal mutations, OL mutations mimic features of somatic mutations in cancers of the CNS, including (1) OL mutational signatures^39^ and (2) the distribution of mutations across the genome, particularly for tumors for which OPCs are believed to be the cell of origin (GBM and oligodendroglioma). These observations emphasize the contribution of mutations that occurred during the OPC stage to the mutation landscape of mature OLs. Furthermore, the differing genomic regions enriched for somatic mutations in OLs versus neurons suggests that cell type-specific mutation distributions may contribute to cell type-specific predispositions to particular pathologies.

## ACKNOWLEDGMENTS

We thank J. Neil and the University of Maryland Brain Bank for helping with the tissue collection. We thank the donors and their families for their invaluable donations for the advancement of science. We thank the Boston Children’s Hospital IDDRC Molecular Genetics Core Facility, supported by NIH award U54HD090255 from the National Institute of Child Health and Human Development.

## AUTHOR CONTRIBUTIONS

J.G, S.B and L.J.L conceived the study; J.G and S.B performed all the experiments; L.J.L led bioinformatic analyses helped by C.L.B H.J and A.G; J.G, and S.B contributed to bioinformatic analysis interpretation; S.B contributed to scRNA-seq data analyses; M.M and Z.Z contributed neuronal PTA data; C.A.W. and P.J.P. directed the research; J.G and L.J.L wrote the manuscript greatly helped by S.B.

## INCLUSION AND DIVERSITY

One or more of the authors of this paper self-identifies as an underrepresented ethnic minority in their field of research or within their geographical location.

## METHODS

### Data and code availability

Newly generated sequencing data for MDA and PTA single-cell WGS of oligodendrocytes, single-cell ATAC-seq, and single-cell RNA-seq are available on dbGaP, accession number `phs001485.v3.p1`. Previously generated scWGS, matched bulk controls, and scRNA-seq data for individual 1465 are also available for download at dbGaP, accession number `phs0014 85.v3.p1`. Scripts used for analyses in this manuscript are available on Zenodo (https://doi.org/10.5281/zenodo.7508802).

### Human tissues and DNA samples

All human tissues were obtained from the NIH NeuroBioBank at the University of Maryland. Frozen post-mortem tissues from 12 neurologically normal individuals were obtained as part of a previous study^4^. Samples were processed according to a standardized protocol (http://medschool.umaryland.edu/btbank/method2.asp) under the supervision of the NIH NeuroBioBank ethics guidelines. Bulk DNA was extracted using the QIAamp DNA Mini kit with RNase A treatment.

### Nuclear sorting and whole-genome amplification

Isolation of single nuclei using fluorescence-activated nuclear sorting (FANS) for NEUN and SOX10 was performed using a modified version of a previously described protocol^53,54^. Briefly, nuclei were prepared by dissecting fresh-frozen human brain tissue previously stored at −80°C, dissolved on ice in chilled nuclear lysis buffer (10mM Tris-HCl, 0.32M Sucrose, 3mM MgAc_2_, 5mM CaCl_2_, 0.1mM EDTA, pH 8, 1mM DTT, 0.1% Triton X-100) using a Dounce homogenizer. Lysates were layered on top of a sucrose cushion buffer (1.8M Sucrose, 3mM MgAc_2_, 10mM Tris-HCl, pH 8, 1mM DTT) and ultra-centrifuged for 1 hour at 30,000rcf. Pellets containing nuclei were resuspended in ice-cold PBS 1X supplemented with 3mM MgCl_2_, then filtered, blocked in PBS 1X supplemented with 3mM MgCl_2_ and 3% Bovine Serum Albumin (blocking solution), and stained with an anti-NEUN antibody (Millipore MAB377) previously used for neuronal nuclei isolation^4,53^, anti-SOX10 antibody (Novus NBP2-59621R), and DAPI. Other antibodies targeting the OL population were also evaluated, KLK6 (Bioss bs-5870R) and CNP (Bioss bs-1000R). Nuclei were washed once with blocking solution, centrifuged at 500rpm for 5 minutes and resuspended again in blocking solution. Nuclear sorting was performed in one nucleus per well in 96-well plates and whole-genome amplification was performed using Primary Template-directed Amplification (PTA) following manufacturer guidelines. Libraries for sequencing were generated using the KAPA HyperPlus kit (Roche) using dual indexes and were sequenced across 5 lanes of Ilumina NovaSeq6000 (2×150bp), targeting 20x coverage (75Gbp)/sample.

### Sorting purity assessment

Three populations were sorted, DAPI, NEUN+, SOX10+/NEUN-, from one 17y individual (Supp. Fig. S1). 10x scRNAseq was used and 6,000-10,000 nuclei from each population were sorted into individual tubes containing RT mastermix, and immediately processed for GEM generation, barcoding, and cDNA amplification, following manufacturer instructions. Each library prep was submitted to paired-end single indexing sequencing on a lane of Illumina HiSeqX to obtain ~50,000 read pairs per nucleus. Analysis of scRNA seq data was performed with Loupe Cell Browser 4.q software provided by 10x genomics. Sorting purity is critical when preforming single-cell whole-genome (scWGS) studies, hence we evaluated purity of a larger number of cells by scRNA-seq. As a control, evaluation of 3,447 DAPI+ sorted nuclei obtained from a mix of grey and white matter (PFC) from a 17yr male identified all the cell-types anticipated for this region and with significant presence of OLs following WM expectation^55^. The purity of 3,739 NEUN+ sorted nuclei was nearly 100% (Supp. Fig. S1A-C) with ~1% (40 out of 3,739 nuclei) presence of a PLP1/MBP/MOG+ OL population, and 0.1% (3 out of 3,739 nuclei) of NOSTRIN+ endothelial cells. NEUN sorted nuclei can be broadly classified into 60% excitatory and 40% inhibitory neurons consistent with recent reports of excitatory/inhibitory ratios^56^. Evaluation of 9,227 SOX10+/NEUN-sorted nuclei confirmed 100% mature OLs purity, with the absence of other cell-type markers expression (Supp. Figure S1A-C). The SOX10+/NEUN-sorted nuclei showed homogenous distribution of classic mature OL-markers such as PLP1, MOG, MALAT1, among others. Although SOX10 is expressed in all stages of OL development, including in OPCs, our strategy consistently recovered only mature OLs.

### 10x scRNA-seq

scRNA-seq was performed using the 10X Genomics Chromium Next GEM Single Cell 3□ Reagent Kit v3.1. Fresh frozen human brain tissue from the prefrontal cortex of individuals UMB1465, UMB4638 and UMB4643 was processed to obtain nuclear pellets. Briefly, tissue was dissociated on ice in chilled nuclear lysis buffer (10 mM Tris-HCl, 0.32 M Sucrose, 3 mM MgAc2, 5 mM CaCl2, 0.1 mM EDTA, pH 8, 1 mM DTT, 0.1% Triton X-100) using a dounce homogenizer. Homogenates were layered on top of a sucrose cushion buffer (1.8 M Sucrose, 3 mM MgAc2, 10 mM Tris-HCl, pH 8, 1 mM DTT) and ultra-centrifuged for 1 hour at 30,000 rcf. Pellets containing nuclei were resuspended in 250 μl ice-cold 1X PBS supplemented with 3 mM MgCl2, 3% Bovine Serum Albumin (BSA) and 0.2 U/μl RNAse inhibitor (Thermo Fisher Scientific ref.10777019), then filtered. After filtering, suspension volume was completed at 1 ml using the same solution, and nuclei were stained with DAPI before sorting to select for intact nuclei. Some of the UMB1465 samples were additionally stained with the following antibodies: two samples with anti-NEUN antibody (Millipore MAB377) for neuron sorting, one sample each for anti-CX43/GJA1 (Novus Biologicals, FAB7737R-1 00UG AF647), anti-SOX9 (Abcam, ab196450 AF488) and anti-GFAP (Millipore, MAB3402 AF647) to enrich for glial cells, and one sample with anti-SOX10 (Novus Biologicals, NBP2-59621 AF647) for oligodendrocyte sorting. 10,000 to 15,000 single nuclei were sorted for each experiment directly in a tube containing the 10X RT mastermix, and immediately processed for gel-bead in emulsion (GEM) generation, barcoding, cDNA amplification and library preparation following manufacturer instructions. Each library preparation was submitted for paired-end single indexing sequencing on Illumina HiSeqX or NovaSeq6000 targeting ~50,000 read pairs per nucleus.

### 10x scRNA-seq data analysis

scRNA-seq data were demultiplexed using `bcl2fastq`. snRNA-seq FASTQ files were then processed using the 10X Genomics `cellranger count` pipeline for gene expression to perform alignment to hg19, barcode counting, UMI counting, and generation of feature-barcode matrices. Cell Ranger filtered count matrices were used for downstream analysis using Seurat 3.0^57^. For each library, we further filtered for cells with > 200 and < 3000 genes and <5% mitochondrial genes, and genes with <10,000 UMI counts and >3 cells. RNA counts were normalized using the LogNormalize method and the 2,000 most highly variable features were identified using the vst method. Data were then scaled by regressing out the percentage of mitochondrial genes. We then performed non-linear dimensional reduction and clustering. To remove doublets from our datasets, we ran DoubletFinder^58^ using optimal parameters as per the paramSweep function. Finally, cell-type identities were assigned to each cluster in the Uniform Manifold Approximation and Projection (UMAP) based on expression of known brain cell-type markers^8^.

### 10x scATAC-seq

Nuclei from 10 individuals (1278, 1465, 4638, 4643, 5087, 5219, 5559, 5817, 5823, 5871) from our aging cohort were obtained from the same brain region as used for single cell whole-genome amplification. Tissue was processed as described in nuclear sorting, and nuclei were re-suspended in diluted nuclei buffer provided by the manufacturer. Nuclei derived from different individuals were processed for transposition separately, before loading to the 10x Chromium Controller for GEM generation, barcoding, and library construction, as per manufacturer instructions. Libraries were submitted for paired-end dual index sequencing on one flow cell of Illumina S2 NovaSeq6000 (100 cycles) to obtain ~50,000 reads per nucleus.

### 10x scATAC-seq data analysis

Sequencing data were demultiplexed using `bcl2fastq` and `mkfastq. cellranger-atac count` v1.1.0 was run separately on the resulting FASTQ files for each scATAC-seq library (one per individual) with default parameters and the vendor-provided hg19 reference. Results from the individual library analyses (Cell Ranger output files `fragments.tsv.gz` and `singlecell.csv` from each library) were then merged by `cellranger-atac aggr --normalize-depth`. scATAC-seq data were analyzed by Signac v1.1.0 and Seurat v3 following the authors’ instructions. Briefly, the merged Cell Ranger output was imported via `Read10X_h5` and `CreateChromatinAssay`; analyzed by `RunTFIDF, FindTopFeatures, RunSVD` and `RunUMAP` with LSI reduction; and integrated with our scRNA-seq to assign cell types via `GeneActivity, FindTransferAnchors` and `TransferData`.

### Single neuron whole genome sequencing data

Sequencing data for 36 PTA-amplified single neurons and matched bulk from the 12 individuals from which OLs were harvested were downloaded from dbGaP accession phs001485.v3.p1. These BAM files were re-analyzed by SCAN2 jointly with OLs as described in *Somatic mutation calling with SCAN2*. For a single additional individual (5171), no PTA data were generated for either neurons or OLs; in this one case, sequencing data for MDA-amplified single neurons from the prefrontal cortex and a matched bulk were downloaded from the same dbGaP accession as the PTA data (see Table S1). SCAN2 somatic SNV calls, indel calls, and genome-wide burden estimates from 15 previously published PTA-amplified single neurons from 5 additional neurotypical individuals (individuals 4976, 5451, 5572, 5666, and 5943) were downloaded from https://github.com/parklab/SCAN2_PTA_paper_2022.

### Somatic mutation calling with SCAN2

SCAN2 v1.1 was run on the 12 individuals from which PTA OLs were collected and individual 5171 (MDA only). First, a cross-sample panel (required for indel calling with SCAN2) was built for the 12 individuals with PTA neurons and OLs by running scan2 config with parameters `--analysis makepanel --gatk=gatk3_joint`; the GRCh37 human reference genome with decoy hs37d5 (`--ref`), dbSNP v147 common (`--dbsnp`) and 1000 Genomes phase 3 SHAPEIT2 phasing panel (`--shapeit-refpanel`) as described in ref. 10; one `--sc-bam` argument for each PTA BAM; and one `--bam` argument for each of the 12 matched bulk BAMs. After generation of the cross-sample panel, SCAN2 v1.1 was run for each of the 12 individuals and individual 5171. For each individual, all PTA OLs (and MDA OLs for individuals 1278, 5871, 5171, and 5657) and PTA neurons (MDA neurons for individual 5171) were analyzed together in a single SCAN2 run. `scan2 config` was run with the parameters `--analysis=call_mutations --gatk=gatk3_joint --abmodel-samples-per-step=20000 --abmodel-refine-steps=4 --abmodel-n-cores=10` and the same GRCh37 reference, dbSNP and phasing panels used above. The cross-sample panel created above was specified via the `--cross-sample-panel` parameter. Finally, SCAN2 mutation signature-based rescue was run in two batches, one for PTA neuron and one for PTA OL calls, using scan2 config `--analysis = rescue --rescue-target-fdr=0.01` followed by `scan2 rescue`. Mutation signature-based rescue was not run on MDA neurons or OLs. For neurons only, VAF-based SCAN2 calls (i.e., excluding signature-rescued SCAN2 calls) from the 15 previously analyzed PTA single neurons (see *Single neuron whole genome sequencing data*) were added to the rescue process via `--add-muts`.

### Total mutation burden estimation and aging models

SCAN2 provides estimates of the total somatic SNV and indel burden in each sample (i.e., the estimated total number of mutations per cell adjusted for sensitivity of mutation calling). These estimates were obtained from each SCAN2 output RDA file by first loading the file in R, then running object@mutburden [[“snv”]]$burden[2]. Indel burdens were recovered by replacing `“snv”` with `“indel”`. To account for variability within and between individuals, mixed-effects linear models were used to estimate the aging-related rates of somatic SNV and indel accumulation. These models were fit by the `R lme4` package using `lmer(genome.burden ~ age*celltype + (1|individual))`, where `celltype` is either `oligo` or `neuron, individual` is one of the 12 individual IDs for which PTA OLs were available (i.e., excluding individual 5171) and age is the numeric age of individual. Confidence intervals were estimated by `confint`. For linear mixed models, statistical tests of significance comparing each coefficient, interaction term and intercept to a null hypothesis of 0 were calculated by the `lmerTest R` package, which uses a *t*-test based on the Satterthwaite approximation. Throughout the text, these *t*-tests are referred to as LMM *t*-tests.

### SnpEff annotation

Both VAF-based and mutation signature-rescued SCAN2 somatic mutation calls were annotated for functional impact via SnpEff version 4.3t using the hg19 database. Reported functional impacts were taken from the first ANN field in the SnpEff annotated VCF. When computing the fraction of mutations with LOW, MODERATE, and HIGH impact (Figure 1D), duplicate and clustered mutations were removed as described in *Somatic mutation enrichment analysis*.

### Somatic mutation enrichment analysis

Enrichment analyses following the methodology described in ref. 10 were carried out to determine whether somatic mutations accumulate preferentially in certain areas of the genome. First, somatic mutations were filtered to remove duplicates and clusters of mutations in single cells. For exact duplicate mutations (i.e., same position and base change or indel), if the duplicate mutations all occur in cells from a single individual (suggesting a clonal mutation), a single representative mutation was retained; otherwise, if the duplicate mutations span multiple individuals (suggesting artifacts), all mutations were discarded. Clusters of mutations, defined as any mutation within 50 bp of another mutation in the same single cell, suggest underlying alignment artifacts or structural variants and were removed before enrichment analysis. Duplicate and clustered mutations were determined separately for MDA and PTA mutation calls. For MDA OLs only, all SCAN2 mutation calls from the 20 OLs from infant brains were additionally filtered prior to duplicate and cluster filtering. These were removed because the mutation burden of young OLs is too small to sufficiently outnumber MDA technical artifacts. The only enrichment analysis using MDA OL calls is the genic region enrichment analysis in Figure S2C.

Next, the genome was divided into regions based on a single covariate (described in more detail below) and the number of somatic mutations in each region *R* was compared to the number of mutations in *R* after the locations of the mutations were randomly permuted across the genome. The permutation process was repeated 10,000 times and the final enrichment value *E_R_* was given by dividing the number of observed somatic mutations in each region *R* by the average number of permuted mutations in *R*. For enrichment analysis of mutational signatures, the above steps were followed, except: (1) somatic mutations called by SCAN2’s mutation signature-based rescue method were not used and (2) rather than counting the number of mutations in each region *R*, the set of mutations in *R* was fit to the reduced COSMIC SBS or ID catalog (see *COSMIC signature catalog activity thresholds reduction*) by non-negative least squares (using lsqnonneg from the pracma R package) and the exposure value for each signature was used in lieu of mutation counts. To obtain two-sided tests of enrichment significance for each region *R*, a permutation test strategy was used. Enrichment values 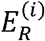 were computed for each of the 10,000 permutations *i* and a *P*-value was derived from the fraction of permutation sets with more extreme enrichment values than the observed mutations. To avoid *P*-values of 0, a minimum of *P* = 0.0001 was enforced, i.e.,

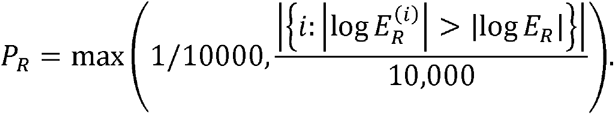

To define regions *R* for enrichment analysis, a subset of the genome with anomalous sequencing depth was first identified and removed from subsequent analyses. The human reference genome GRCh37 with decoy sequences hs37d5 was divided into non-overlapping windows of 100 bp and the average sequencing depth across all PTA neurons and OLs as output by SCAN2 (file path: path/to/scan2_output/depth_profile/joint_depth_matrix.tab.gz) was computed for each 100 bp window. Windows with low average depth (<6 reads averaged across all PTA cells) or excessive average depth (in the top 2.5% of average depth) were classified as anomalous. Genomic regions were next derived from non-quantitative genomic covariates (genes, Figure S2D; exons, introns, and intergenic spaces, Figure 1C; ChromHMM classes, Figure 4F; and cell type-specific promoters and enhancers, Figure 4G) and quantitative covariates (GTEx transcription levels, Figure S3A; scRNA-seq transcription levels, scATAC-seq accessibility, RepliSeq replication timing and histone mark levels, Figures 4B,D-G). For non-quantitative covariates, regions *R* were defined by the union of genomic intervals for each unique covariate state (e.g., all exons or all regions annotated as ChromHMM state 1) and anomalous windows were subtracted from these unions. For quantitative covariates, the genome was first tiled with non-overlapping 1 kbp windows (corresponding to 10 100 bp windows from the anomalous sequencing depth analysis). 1 kbp windows containing >2 anomalous sequencing depth windows were discarded. For each remaining 1 kbp window *i*, a single quantitative value *V_i_* was derived for each covariate in a covariate-dependent manner (described in detail for each covariate below). The distribution of values *V_i_* were then discretized into n = 10 (for enrichment analysis of total mutation burden) or n = 3 quantiles (for enrichment analysis of mutation signatures) and each window *i* was assigned its quantile rank *Q_i_*. Finally, a region *R_Q_* was defined for each quantile *Q* = 1…n by taking the union of windows with rank *Q*. Genomic regions and the resulting enrichment analyses were always performed using one covariate at a time.

### Genomic covariates for enrichment analysis

#### Gene regions

GENCODE genes version 26 was downloaded from https://ftp.ebi.ac.uk/pub/databases/gencode/Gencode_human/release_26/GRCh37_mapping/gencode.v26lift37.annotation.gtf.gz. A single region was defined for each protein coding gene using GTEx’s transcript collapse script (https://raw.githubusercontent.com/broadinstitute/gtex-pipeline/master/gene_model/collapse_annotation.py) and used for the per-gene enrichment analysis in Figure S2D. Exonic, intronic and intergenic regions were then derived from the GENCODE GTF records for the same set of genes. When multiple records overlapped a single locus, status was prioritized as follows: CDS > UTR > exon > intron > upstream/downstream > intergenic. Finally, loci classified as CDS, UTR or exon were classified as exonic, intron as intronic, and all other classes as intergenic.

#### ChromHMM chromatin states

15-state ChromHMM annotations were downloaded for epigenome ID E073 (dorsolateral prefrontal cortex) from https://egg2.wustl.edu/roadmap/data/byFileType/chromhmmSegmentations/ChmmModels/coreMarks/jointModel/final/E073_15_coreMarks_mnemonics.bed.gz.

#### Promoters and enhancers for flow sorted brain cell types

Active promoter and enhancer elements were extracted from Supplementary Table 5 of ref. 58 available at https://www.science.org/doi/suppl/10.1126/science.aay0793/suppl_file/aay0793-nott-table-s5.xlsx. Duplicate lines in these tables were removed prior to analysis.

#### GTEx transcription levels

Median gene expression levels from 54 tissue types were downloaded from the GTEx project at https://storage.googleapis.com/gtex_analysis_v8/rna_seq_data/GTEx_Analysis_2017-06-05_v8_RNASeQCv1.1.9_gene_median_tpm.gct.gz. For each tissue type, the median transcription level of each gene *G* was mapped to the genome by applying it over *G*’s collapsed transcript (see *Gene regions* above). Only 1 kbp genomic windows with >=80% coverage by gene transcripts were retained. If multiple genes overlapped the same window, the maximum TPM value was used for the window.

#### scRNA-seq transcription levels

Cell type-annotated gene-expression matrices for each scRNA-seq library were concatenated column-wise and average expression levels for each gene were calculated for each cell type. Gene names were then matched to the GTEx gene model and transcription levels for each cell type were mapped to the genome as described above for GTEx transcription levels.

#### scATAC-seq accessibility

ATAC-seq transposition events output by cellranger-atac (file: fragments.tsv.gz) were first separated by cell type (see *10x scATAC-seq data analysis*) and then converted to BED format. The BED file of fragments for each cell type was then converted to bedGraph format using bedtools genomecov -bga and finally to bigWig format by bedGraphToBigWig. The bigWig signal files were then mapped to the 1 kbp genome tiles described in *Somatic mutation enrichment analysis* by bigWigAverageOverBed.

#### Replication timing

WaveSignal RepliSeq bigWigs were downloaded from http://hgdownload.cse.ucsc.edu/goldenpath/hg19/encodeDCC/wgEncodeUwRepliSeq/wgEncodeUwRepliseq{cell_line}WaeSignalRepl.bigWig for 15 cell_lines (all available at the time of writing): BG02ES, BJ, GM06990, GM12801, GM12812, GM12813, GM12878, HUVEC, HeLa-S3, HepG2, IMR90, K562, MCF-7, NHEK and SK-N-SH. The bigWig signal files were then mapped to the 1 kbp genome tiles described in *Somatic mutation enrichment analysis* by bigWigAverageOverBed; quantile values were then reversed so that *Q* = 1 corresponded to the earliest replication timing quantile.

#### Histone marks

bigWig signal files representing ChIP-seq fold-change versus a no-IP control were downloaded for 13 Roadmap reference epigenomes annotated as brain tissue from https://egg2.wustl.edu/roadmap/data/byFileType/signal/consolidated/macs2signal/foldChange/{epigenome_ID}-{histone_mark}.fc.signal.bigwig for 7 histone_mark values H3K27ac, H3K27me3, H3K36me3, H3K4me1, H3K4me3, H3K9ac and H3K9me3. The bigWig signal files were then mapped to the 1 kbp genome tiles described in *Somatic mutation enrichment analysis* by bigWigAverageOverBed.

### COSMIC signature catalog activity thresholds and reduction

To reduce the possibility of overfitting by signatures that may not be active in our cell types, all signature fits were conducted on a subset of the COSMIC v3.1 SBS and ID catalogs. The catalogs were reduced by several heuristics designed to remove signatures with low likelihood of activity in our data. The first heuristic is a step-forward procedure in which each step *i* determines the single COSMIC signature that minimizes the residual (resid.norm) of the non-negative least-squares fit (lsqnonneg) to the (signature channel) *×* (sample) input matrix compared to the signatures present in step (*i* – 1). At each step, this reduction in residual is compared to the residual using 0 signatures (i.e., the norm of the input matrix) to determine the percent reduction of the signature selected at each step *i*. The second heuristic is designed to find evidence of age-related accumulation of each signature. First, the (signature channel) × (sample) input matrix is fit by lsqnonneg to the entire COSMIC SBS or ID catalog to determine exposure levels *E_i,j_* for each sample *i* and signature *j*. For each signature *j*. a linear model *E*, ~ age_*i*_ is fit and the significance of the test for slope=0 (*t*-test) is recorded. Finally, the significance values for all signatures tested are corrected for multiple hypothesis testing by `R’s p.adjust` method using Holm’s correction. The third heuristic measures the percent contribution *Pj* of each signature *j* to the total spectrum; i.e., *P_j_* = ∑*_i_ E_i,j_*/∑*_i,j_ E_i,j_*. These three heuristics are computed separately for neurons and OLs using VAF-based SCAN2 somatic mutation calls (to avoid signature biases of signature-rescued SCAN2 calls). For each set of heuristics, signature *j* is retained if either: heuristic (1) exceeds a 50% reduction in the residual; or if heuristic (2) < 0.01 after multiple hypothesis correction and heuristic (3) exceeds 1% of the total spectrum. The final set of retained signatures is the union of signatures retained in either neurons or OLs.

### Analysis of related oligodendrocyte pairs

Shared somatic SNVs were determined for each pair of single cells within each individual. A shared sSNV was defined as a SCAN2 call present in at least one of the two cells and for which 2 or more mutation supporting reads appear in the other cell. Private mutations were SCAN2 calls that: (1) did not meet the shared mutation criteria and (2) were supported by 0 reads and a total depth of 6 or greater in the paired cell. These heuristics identified two pairs of oligodendrocytes with exceptionally high numbers of shared sSNVs (5559-Oligo-5 and 5559-Oligo-8, PTA-amplified OLs from individual 5559; 5657-GliaLC-4-F11 and GliaLC-4-G10, MDA-amplified OLs from individual 5657).

Because the pair of OLs from individual 5657 were MDA-amplified, it was necessary to remove artifacts associated with MDA, which follow a specific SBS signature termed Signature B^10^. For signature analysis, removal of this signature was achieved by adding Signature B to the reduced COSMIC catalog before fitting by lsqnonneg and then discarding exposures to Signature B. Although PTA induces a much lower artifact burden than MDA, a similar correction was performed to remove PTA artifacts from the spectra presented in Figure 2D,E: the PTA artifact signature was added to the reduced COSMIC catalog, mutations were fit by lsqnonneg and finally the reconstructed spectrum was created by summing COSMIC signatures, weighted by their exposures, and excluding the PTA artifact weights.

To estimate time to the most recent common ancestor (MRCA) split, several corrections to account for sensitivity and shared vs. private misclassification were required. To quantify shared vs. private misclassifications, we applied the shared and private criteria for somatic SNVs to germline heterozygous SNPs (hSNPs) known to be present in the individual, which should always be shared between single cells from the same individual. The number of shared sSNVs was then multiplied by (1 + fraction of hSNPs classified as private). The number of private sSNVs was then reduced by the amount added to the shared sSNV count. The opposite error, a private sSNV classified as shared, should occur rarely since it requires a random artifact to intersect with a true mutation; we thus assumed this rate to be approximately 0. For the pair of MDA OLs, the number of private sSNVs was reduced by the fraction of the private sSNV spectrum attributed to the MDA artifact Signature B after fitting to the COSMIC SBS catalog with Signature B. SCAN2’s estimate of the total mutation burden (which accounts for mutation calling sensitivity) was also reduced by the Signature B exposure fraction and SCAN2’s calling sensitivity re-estimated by *S* = (# SCAN2 calls) / adjusted total burden. The final corrected shared sSNV and private sSNV counts reported in Figure 2D were computed by dividing by *S*. The time to MRCA was estimated by counting mutations forward from the initial zygote (i.e., using the number of corrected shared sSNVs) and backward (i.e., using the number of corrected private sSNVs). In each case, the corrected number of mutations was converted to a time in years by first subtracting the intercept of our OL sSNV aging model and then dividing by the slope of the OL aging model.

### Cancer mutation density

PCAWG cancer somatic SNV and indel mutation catalogs were obtained in MAF format from the ICGC portal (https://dcc.icgc.org/releases/PCAWG/consensus_snv_indel). Hypermutated tumors within each tumor type were identified by Tukey’s method and mutations from these tumors were removed. Next, cancer mutations for each sample were mapped to the 100 bp windows used for detecting anomalous sequencing depth (described in *Somatic mutation enrichment analysis*) and the count of mutations in each window was then normalized by the total number of mutations in that sample. Finally, a track representing the mutation density for each tumor type was created by summing normalized window counts across samples from the same tumor type and written in bigWig format via `rtracklayer’s export.bw` function.

Because 100 bp or 1 kbp windows contain too few mutations for meaningful correlation analyses with our neuron and OL somatic mutations, the per-tumor bigWig signal files with 100 bp resolution were mapped to a non-overlapping 1 Mbp genome tiling by `bigWigAverageOverBed`. 1 Mbp windows with anomalous sequencing depth were then removed following the same requirements for the 1 kbp tiling windows described in *Somatic mutation enrichment analysis*. Total neuron and OL mutation counts were also determined over these 1 Mbp windows and correlations between somatic mutation density in OLs, neurons and each tumor type were shown in Figure 6A. A similar downscaling of signals from 1 kbp resolution to 1 Mbp resolution via `bigWigAverageOverBed` was required for comparing cancer mutation densities to scRNA-seq and scATAC-seq signals.

### Cancer gene odds ratio analysis

Since the sizes of individual genes are usually too small for detecting mutation enrichment or depletion with our catalog of somatic mutations from OLs and neurons, we created a larger genomic region by considering sets of genes. PCAWG mutations (without hypermutated samples, as described in *Cancer mutation density*) were mapped to genes by SnpEff as described above. For each tumor type, a list of genes ordered by number of somatic mutations intersecting the gene was constructed. For each tumor type, 50 progressively larger genomic regions corresponding to the union of the top 10, 20,… 500 genes were created. In each region *R*, the rates of OL mutations and neuron mutations impacting the region were compared using the odds ratio

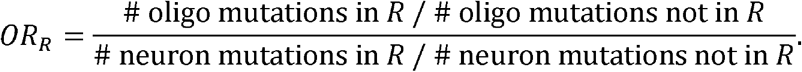

Thus, *OR_R_* > 1 implies a preference for OL mutations in the genes represented in *R* and *OR_R_* < 1 implies a preference for neuronal mutations in *R*. Figure 6D presents odds ratios for the top 100 genes in each tumor type and the full analysis is presented in Figure S5.

Supplementary Information is available for this paper.

Correspondence and requests for materials should be addressed to CAW and PJP.

