## Supplementary figures for "Contrasting patterns of somatic mutations in neurons and glia reveal differential predisposition to disease in the aging human brain"

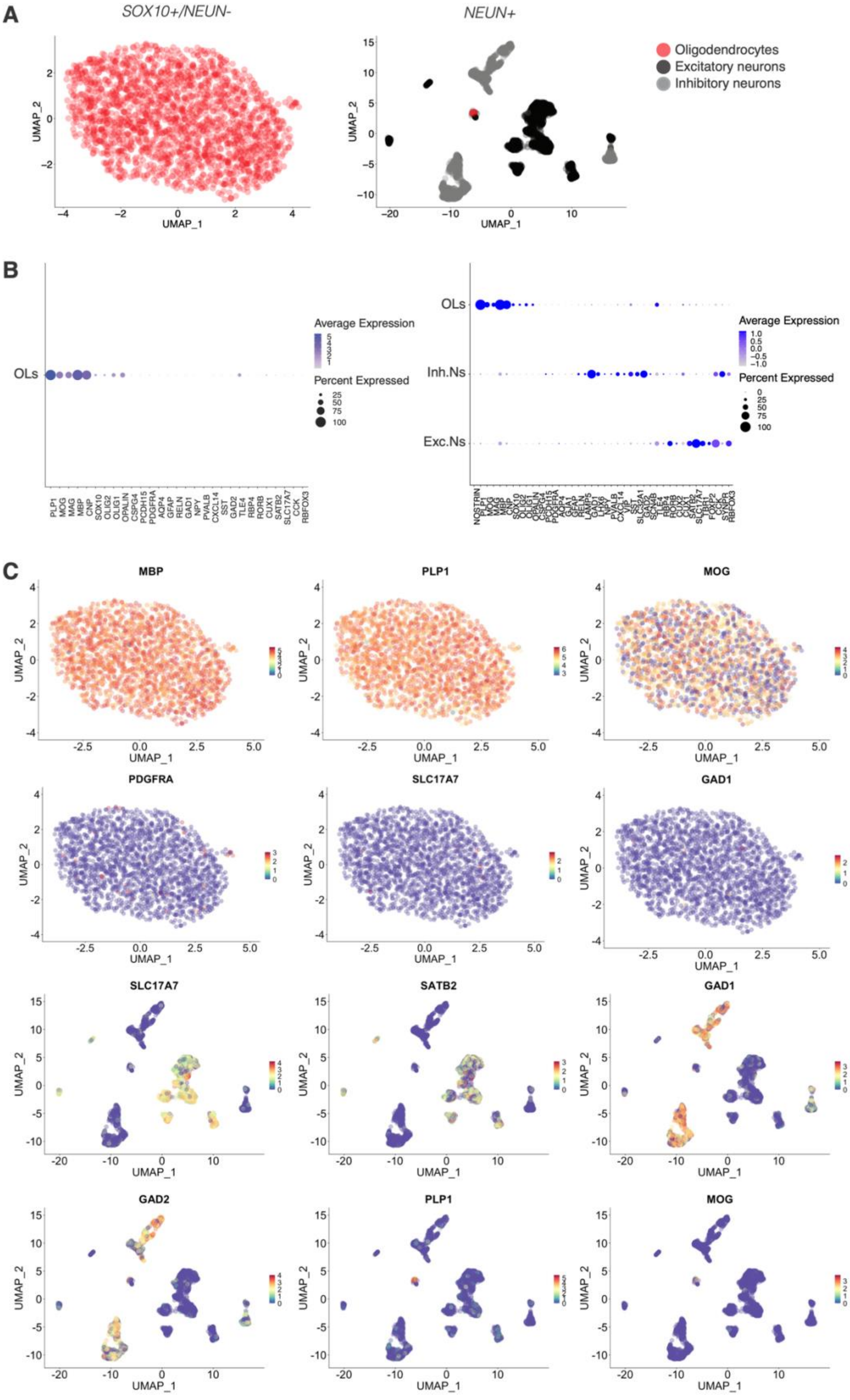
A
SOX10+/NEUN-
NEUN+
Oligodendrocytes
Excitatory neurons
Inhibitory neurons
UMAP_1
UMAP_2
B
OLs
Inh.Ns
Exc.Ns
Average Expression
Percent Expressed
PLP1 MOG MAG MBP CNP SOX10 OLIG2 OLIG1 OPALIN CSPG4 PCDH15 PDGFRA AQP4 GFAP RELN GAD1 NPY PVALB CXCL14 SST GAD2 TLE4 RBP4 RORB CUX1 SATB2 SLC17A7 CCK RBFOX3
C
MBP
PLP1
MOG
PDGFRA
SLC17A7
GAD1
SATB2
GAD2

**Figure S1. Purity of oligodendrocyte and neuron sorting assessed by scRNA-seq, related to Figure 1 and Methods.**
(**A**) UMAP plots demonstrating clusters obtained for SOX10+/NEUN- and NEUN+ FANS experiments. The SOX10+/NEUN- sort gave approximately 100% OLs (left), while NEUN+ sorts contained almost exclusively neurons, both excitatory and inhibitory, and an extremely small percentage (1%) of OLs (right).
(**B**) Dot plot showing expression of cell-type marker genes in the two sorts.
(**C**) Feature plots showing expression of cell type-specific markers. SOX10+/NEUN- sorted nuclei (top two rows) express the OL markers *MBP*, *PLP1* and *MOG* but show no expression of the OPC marker PDGFRA, the excitatory neuron marker *SLC17A7* or the inhibitory neuron *GAD1* markers. NEUN+ sorted nuclei (bottom rows) express *SLC17A7* as well as another excitatory neuron maker *SATB2*, and the inhibitory neuron markers *GAD1* and *GAD2*, but do not express OL markers *PLP1* and *MOG*. Units of transcription are scaled TPM (see **Methods**).

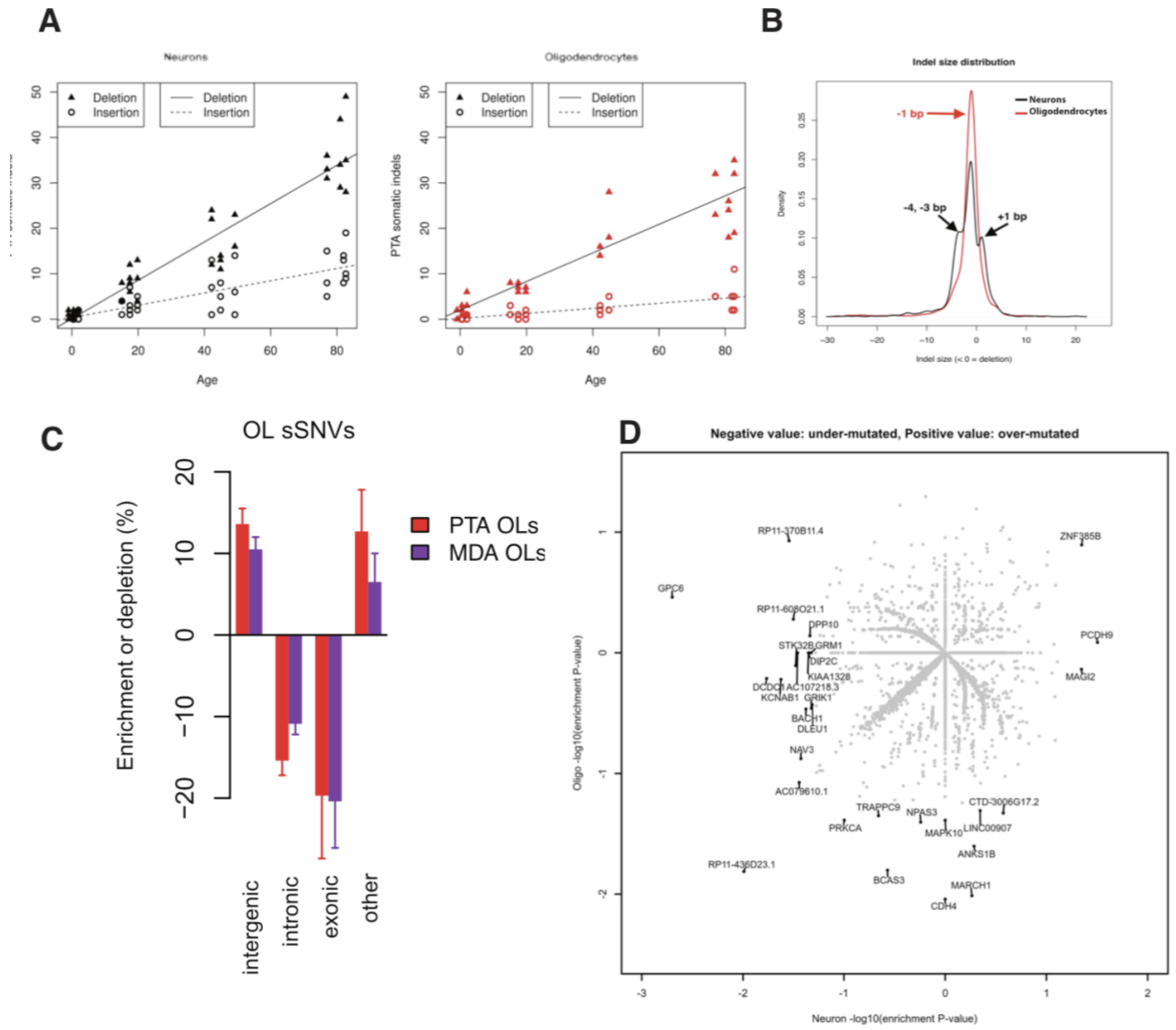


**Figure S2. Somatic indel characteristics, comparison of MDA and PTA sSNV distribution in genic regions and enrichment analysis of individual genes, related to Figure 1.**

**(A)** Separate aging trend lines for insertions and deletions.

**(B)** Distribution of somatic indel sizes. Positive sizes indicate insertions, negative sizes indicate deletions.

**(C)** Comparison of PTA OLs and elderly MDA OLs (*n* = 20 OLs). The mutation burden of elderly (~80 years of age) OLs is high relative to the rate of technical MDA artifacts, allowing enrichment analysis with respect to large genomic regions.

**(D)** Enrichment analysis of individual genes; each point represents a single gene. x- and y-axis values represent significance of enrichment, not enrichment level; significance is signed to represent enrichment (positive significance values) or depletion (negative significance values) of mutations in each gene. Significance values shown are not corrected for multiple hypothesis testing; when corrected, no *P*-values are significant, reflecting the low power to detect enrichment in genes, which are small compared to all other genomic regions considered in this study.

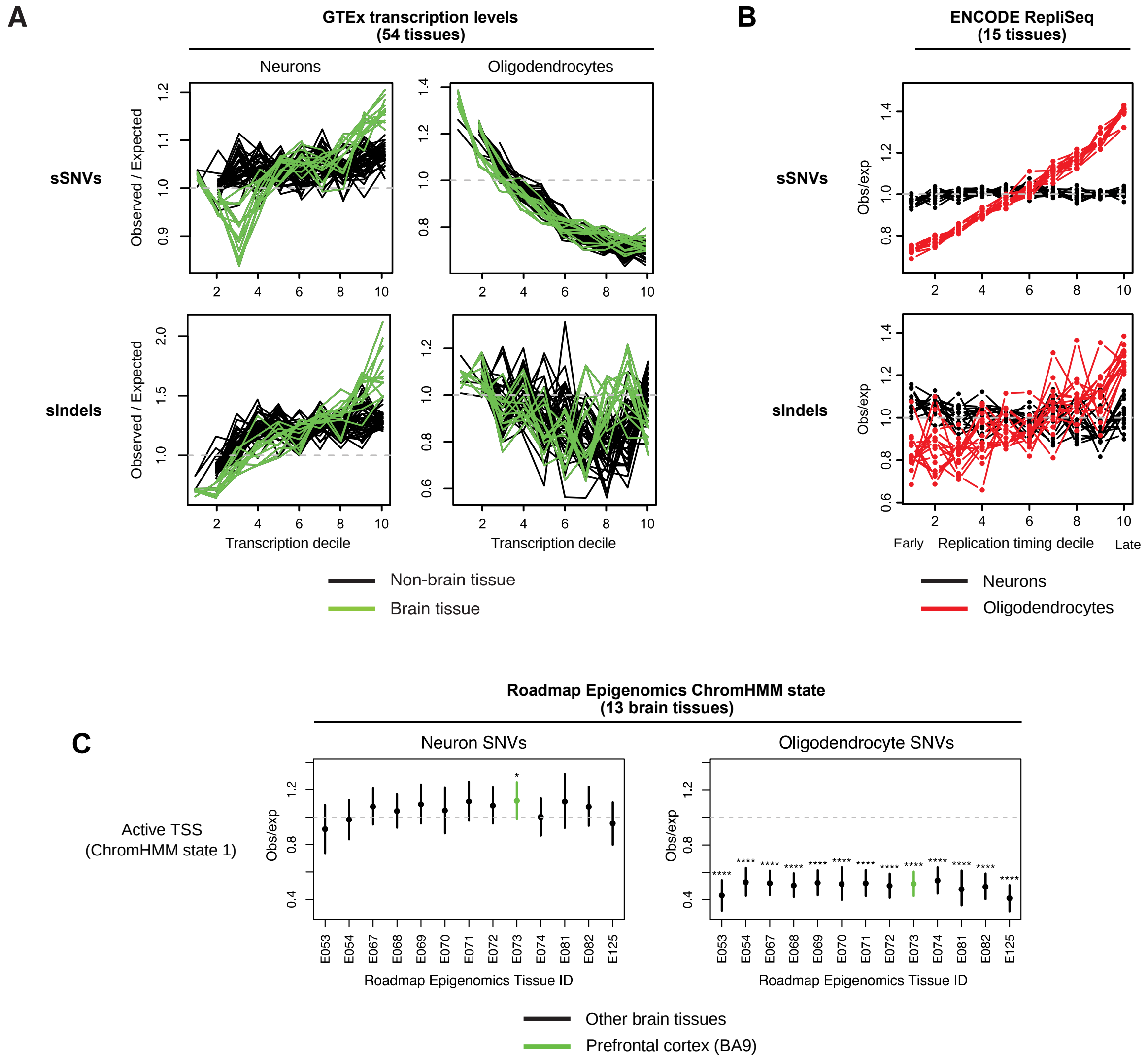


**Figure S3. Somatic mutation enrichment analysis of genomic covariates from diverse tissue types, related to Figure 4.**

**(A)** Comparison of somatic mutation density against publicly available bulk RNA-seq transcription data from GTEx. Each line represents one of 54 tissues in GTEx; green lines, brain tissues; black lines, all other tissue types.

**(B)** Comparison of somatic mutation burden with 15 ENCODE cell lines for which RepliSeq data is publicly available. The averages of these lines (for each of neurons and OLs, separately) are presented in Figure 4E.

(**C**) Enrichment analysis of active promoter regions (ChromHMM state 1_TssA) in 13 brain reference epigenomes from the Roadmap Epigenomics Project. Prefrontal cortex (BA9), colored green, matches the tissue from which neurons and OLs were obtained in this study. * - enrichment *P*-value < 0.1 ; **** - enrichment *P*-value < $10^{-4}$.

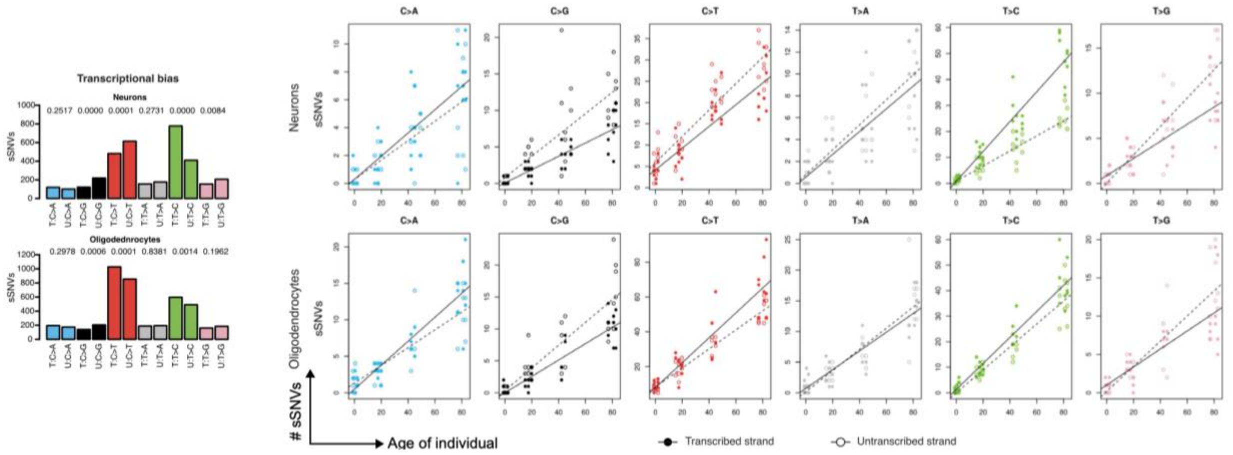


**Figure S4. Transcribed-strand bias of somatic SNVs, related to Figure 5B.**
Levels of transcribed-strand bias vs. age. (left) Aggregate transcribed strand bias across all neurons (top left) and OLs (bottom left); *P*-values above each pair of bars: for each of the six possible single base substitutions, Wilcoxon rank-sum test between sample-specific counts of transcribed and untranscribed mutations. (right) Transcribed-strand bias plotted separately for each single neuron (top right) and OL (bottom right).

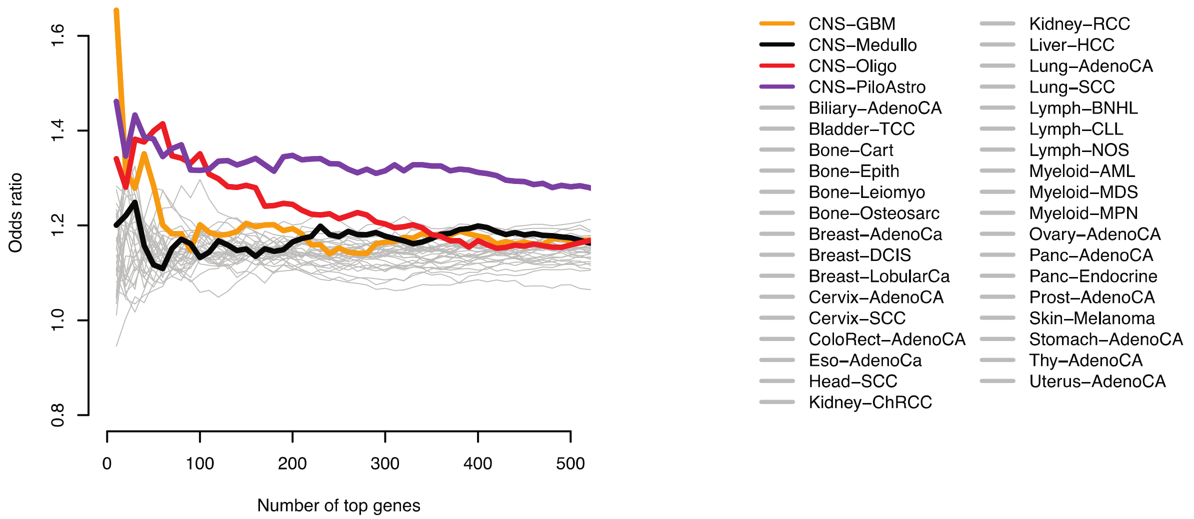


**Figure S5. Oligodendrocyte sSNVs remain frequently enriched in brain cancer genes using a wide range of *n* cutoffs, related to Figure 6.**
Same analysis presented in Figure 6D using several cutoffs to show that choice of top *n* cutoff for the most frequently mutated genes in each tumor type does not affect our conclusions. The analysis in Figure 6D corresponds to $n$ = 100 (x-axis value).
